# Efficient, robust, and versatile fluctuation data analysis using MLE MUtation Rate calculator (mlemur)

**DOI:** 10.1101/2023.01.05.522850

**Authors:** Krystian Łazowski

## Abstract

Almost 80 years since the publication of the seminal work by Salvador Luria and Max Delbrück, the fluctuation assay has remained an important tool for analysing the levels of mutagenesis in microbial populations. The mutant counts originating from some average number of mutations are usually assumed to obey the Luria–Delbrück distribution. While several tools for estimating mutation rates are available, they sometimes lack accuracy or versatility under non-standard conditions. In this work we developed extensions to the Luria– Delbrück protocol to account for phenotypic lag and cellular death with either perfect or partial plating. Hence, our novel MLE MUtation Rate calculator, or mlemur, is the first tool that provides a user-friendly graphical interface allowing the researchers to model their data with consideration for partial plating, differential growth of mutants and non-mutants, phenotypic lag, cellular death, variability of the final number of cells, post-exponential-phase mutations, and the size of inoculum. Additionally, mlemur allows the users to incorporate most of these special conditions at the same time to obtain highly accurate estimates of mutation rates and *p* values, confidence intervals for an arbitrary function of data (such as fold) and perform power analysis and sample size determination for the likelihood ratio test. We assess the accuracy of point and interval estimates produced by mlemur against historical and simulated fluctuation experiments. We believe both mlemur and the analyses in this work might be of great help when evaluating fluctuation experiments and increase the awareness of the limitations of the widely-used Lea–Coulson formulation of the Luria-Delbrück distribution in the more realistic biological contexts.

**Author Summary:** Despite many recent developments in the department of fluctuation data analysis, geneticists are often limited to either a very strict classical Luria–Delbrück protocol, or possibly a modified one with relaxation of just one assumption (e.g., differential fitness of mutants and wild-type cells). In some cases, such as partial plating, researchers use historical methods whose accuracy strictly depends on the conditions of the experiment and thus are not optimal in a wide range of parameters. A novel tool for fluctuation data analysis, mlemur (MLE MUtation Rate calculator), alleviates these problems and provides new extensions that allow to account for phenotypic lag and cellular death. Additionally, we find that the failure to properly account for these additional parameters might lead to inaccurate point and interval estimates of the mutation rate. Our results underline the importance of careful examination of the fluctuation assay to ensure that the chosen statistical model reflects the biological and technical conditions of the experiment.

## 1. Introduction

### 1.1. The concept of measuring mutagenesis in microorganisms

In microbial genetic studies, researchers are often compelled to estimate the mutation rate, that is, the pace at which mutations are accumulated within the genome under certain conditions in a given genetic background and specific organism. Genetic mutations can arise, i.a., during DNA replication, DNA repair, or upon exposure to certain endo- and exogenous agents that chemically change identity of the nitrogenous base within the nucleotide [1,2]. Whatever the mechanism, mutation rate in a specific genetic background can often be informative of the underlying biological processes that affect the fidelity of DNA replication or the effectiveness of DNA repair.

Numerous assays have been developed to score mutagenesis. A modern approach to this problem is deep sequencing of genomic DNA of colonies that underwent multiple passages and accumulated mutations after hundreds of generations [3]. This method is very precise because it is more robust against selection of only certain groups of mutations (for example, silent mutations can be observed). Moreover, it allows to analyse not only the rate, but also the specificity of occurring mutations. On the other hand, mutation accumulation assays are time-consuming, laborious, and expensive. Other existing methods of measuring the mutator phenotype are based on counting the number of mutant cells that acquired a selectable mutation in a certain reporter gene, either before, during, or after the culture/colony growth. These mutants can be sorted and counted in flow cytometry (e.g., mutations in the gene encoding GFP can be scored with the usage of FACS [4]), but most usually are plated, allowing mutant colonies to be counted on solid medium containing a selective agent (usually an antibiotic, a carbon source, or an amino acid).

These kinds of experiments usually start with a number of small parallel cultures of a given microorganism, grown to saturation under specified conditions. After growth, the whole, or a portion of, culture is plated onto the solid medium containing some selective agent. A small portion (usually dilution) of all or only selected cultures is plated on a non-selective medium to estimate the size of the population in each sister culture. After that, the plates are incubated until visible colonies are formed [5].

Since with this approach mutations are not observed directly, but rather by their effect on cell physiology (i.e., the ability to grow on selective medium), in order to infer conclusions about the mutation rate from the number of mutants on the plate, a certain statistical model must be applied. This is caused by the fact that the population of mutant cells (as well as non-mutants) grows exponentially, and the final number of mutants in the culture depends not only on the mutation rate, but also on the time the mutation happened and how many divisions (generations) the mutant cell underwent afterwards (Figure 1A). This type of experiment is the oldest approach to mutation rate scoring and historically has been called a fluctuation assay. The first fluctuation assay was described by Salvador Luria and Max Delbrück in 1943 and eventually inspired generations of biostatisticians and microbiologists to pursuit novel, more precise and accurate methods of measuring mutagenesis in living cells. To this day, fluctuation assay is a very commonly used tool to estimate mutation rates in both bacterial and eukaryotic microorganisms, as well as some human cell lines.

**Figure 1.**
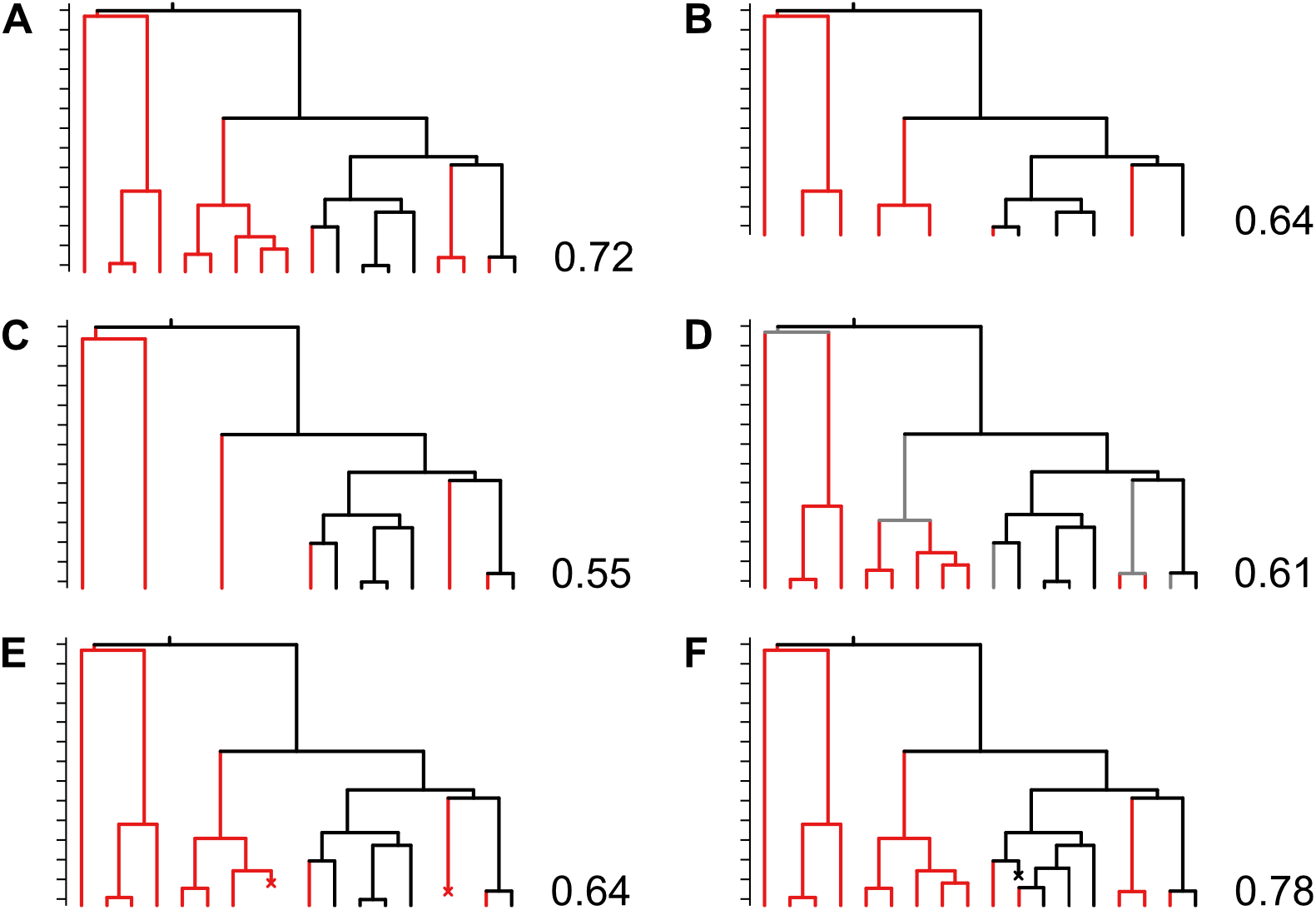
The graphs depicting the interplay between the number of mutations and the proportion of mutant cells in the culture under different conditions. Data were simulated using draw.clone function from R package flan assuming exponential cell lifetimes, *t* = 3, and mutation probability = 0.5. The proportion of mutants in the culture is presented next to each graph. Black lines – wild-type cells. Red lines – mutant cells. Gray lines – mutant cells that have not yet expressed mutant phenotype. Tilted cross – cell death. **(A)** The observed number of mutant cells depends on the stage at which mutation occurred. While most mutations are expected to occur late during exponential growth of the culture (because most cell division happen at this time), the “early” mutations lead to the presence of so-called “jackpot” cultures with exceptionally high number of mutant colonies. **(B)** If the growth of the culture was interrupted at an earlier moment, the produced mutant frequency would be lower (0.64 *vs*. 0.72) despite constant mutation rate. **(C)** The observed mutant colony count depends on the growth rate of the mutant cells in relation to the growth rate of the non-mutant cells. Here, due to the mutant growth rate equal to half the growth rate of the non-mutant cells, the mutant frequency is 0.55. **(D)** The presence of a phenotypic lag decreases the observed mutant colony count. Here, with a phenotypic lag equal one generation, the two last mutations produce cells that fail to grow on selective medium because the mutant phenotype is not expressed at the time of plating (proportion of mutants 0.61). **(E)** Death of a mutant cell decreases the mutant count (mutant frequency 0.64). **(F)** However, death of a non-mutant cell increases the number of cellular divisions needed to reach a given culture size, inflating mutant count (mutant frequency 0.78).

Mutations occurring in an unperturbedly growing microbial culture are typically modelled using the Luria–Delbrück distribution [6], sometimes also called Lea–Coulson distribution to acknowledge their big contribution to the topic [7]. Because there are no explicit expressions for the mean and standard deviation of the Luria–Delbrück probability distribution, it is a common practice to use approximate methods such as Jones method of the median, Lea– Coulson method of the median, Drake formula, and methods based on the mean number of mutants. Since there is a finite probability that a selectable mutation will occur quite early during culture growth, giving rise to an exceptionally big number of mutants on the plate (so-called “jackpot” culture), the mutant distribution is heavily tailed, and therefore methods of mutation rate calculation based on the mean number of mutant cells per plate are significantly biased; so are, although to a somewhat lesser extent and particularly with small samples, methods based on the median. Sometimes scientists are confined to report only mutant frequency (often incorrectly called mutation frequency), that being the average number of mutants observed per some number of cells. Mutant frequency, however, is not a biological property and strongly depends on the number of generations the culture underwent. Because of that, mutant frequencies measured by different protocols are not comparable and are not informative with regard to the number of mutations that led to observed number of mutants (Figure 1A, B).

### 1.2. The Luria–Delbrück distribution and the fluctuation protocol

The Luria–Delbrück mutant distribution arises as a combination of several simple ideas. The basis of the derivation that is easy to follow was developed by Stewart et al. and we will present it here [8,9]. First, we assume that the non-mutant cells grow deterministically (without randomness) but non-synchronously, according to a well–known equation for exponential growth:

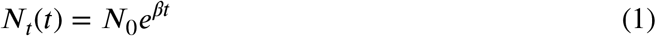

where *N*_0_ is the size of the inoculum, *β* is the growth rate of the cells, and *t* is the time when we take the measurement of *N_t_*. In the context of mutation rate estimation with the usage of the fluctuation protocol, the only time that is of interest is usually the time when the culture stopped growing, so we can equate *t* with the time when we interrupted culture growth and *N_t_* with the size of the culture at this time. The exact value of *t* and *β* is not important because they are implicitly but unequivocally described by *N_t_*. If we define the mutation rate *μ* as the chance of mutation per cell division (the number of cell divisions is obviously *N_t_ – N_0_*, but since the typical size of the inoculum is of the order of magnitude 10^2^–10^3^ and typical final culture size has the order of magnitude 10^8^–10^9^, in most cases we can ignore *N_0_*), and further assume that it is constant over time (i.e., independent of time), then

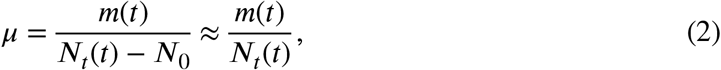

with *m(t)* being the number of mutations that happened in the culture up to the time *t*. From this it follows that the number of mutations is given by *μN*_0_*e^βt^*.

As a side note, the unitless mutation rate per cell division defined in (2) does not depend on the growth rate of the cells. It is commonly assumed that this form of mutation rate does not change when growth rate changes. The same assumption is used throughout this paper. Mutation rate per unit of time can be obtained by multiplying (2) by the independently estimated growth rate of the strain, although it is rarely reported. Multiplication by other factors (2 or logarithm of 2) which were proposed Luria & Delbrück [6] and Armitage [10] has been discouraged [11].

If we consider a very short period of time *dt*, during which there are *dN* cell divisions, then the chance of mutation occurring between *t* and *t* + *dt* is given by the differential

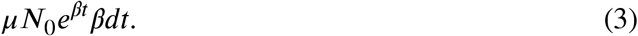

Now, let us say that at some point during culture growth a mutation arose: one wild-type cell divided into one wild-type cell and one mutant cell. If this mutation spanned at some time *τ* before *t*, then the mutant cell has time *t*–*τ* to proliferate. We could model the dynamics of the mutant cells using the same equation (1), as this was done originally by Luria and Delbrück, but their growth is frequently taken as random. A stochastic process similar to the deterministic exponential growth process is called Yule pure birth process, in which the chance of a single mutant cell growing to a clone of size *n* between the time mutation happened *τ* and the time the culture stopped growing *t* is given by (equation (8.15) in Bailey 1964 [12])

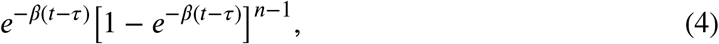

which is also an expression for probability mass function of geometric distribution with *p* = *e^-β(t-τ)^*. In the Yule process the lifetimes of individual cells are exponentially distributed with mean equal *β*. For large *t* both deterministic and stochastic models give asymptotically equivalent results [13]. If we now extend our considerations to the whole time of the culture growth from the beginning to the end at time *t*, the number of mutations giving rise to *n* colonies is given by the integral of the product of equations (3) and (4) over the whole time of culture growth, that is,

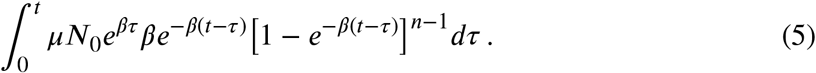

At this moment it is worth to point out that in (2) we conveniently overlook the fact that in a real-world scenario we replace *N_t_* with the average culture size, which contains both wild-type and mutant cells, and the true number of wild-type cell divisions is in fact smaller. Because the estimate of *N_t_* usually has only 3 significant digits, the additional inaccuracy introduced by ignoring mutant cells (similarly to neglecting inoculum) when calculating the number of divisions is miniscule, so *N_t_* + *n* ≈ *N_t_*. However, for this reason mutant cell population relative to *N_t_* should be kept small so that the error does not grow out of control.

The Luria–Delbrück mutant distribution is, of course, merely an approximation of a complex biological process. As a consequence of the model, a classic fluctuation assay has a set of assumptions: cells (a) grow exponentially and (b) independently from each other with (c) no death events (d) from an inoculum that is miniscule in size compared to the final culture; mutations occur (e) at the moment of cell division (f) at a low and (g) constant rate from the time the culture growth started until it stopped, (h) are not influenced by previous mutations, (i) do not revert, (j) do not occur after culture growth has stopped (either in the stationary phase or after plating), are (k) immediately expressed, and (l) result in creation of only one mutant cell (this refers to the moment of mutation and should not be confused with further proliferation of this single initial mutant cell); mutant cells (m) have life-times that obey exponential distribution, (n) constitute a small portion of the whole culture, (o) have comparable fitness to non-mutant cells, and (p) are always detected; (q) parallel cultures are homogenous in terms of volume and size.

While most of the conditions imposed by the Luria–Delbrück model can be easily met, many commonly used fluctuation protocols do not strictly meet its criteria. The requirement that the cells are in the exponential phase of growth is most commonly violated. The cultures are usually grown to saturation (frequently overnight), and therefore the cells undergo growth deceleration and eventually enter the stationary phase. How this affects the presumptive constancy of mutation rate during growth is not well understood; it is known, however, that some unicellular organisms entering the stationary phase activate SOS-induced DNA polymerases and start to accumulate mutations that increase their fitness under conditions of nutrient depletion [14–16].

In the above example the potential conflict between assumptions and protocol can be circumvented by interrupting the culture growth when cells are still in the exponential phase. However, there are many situations where deviations from the fluctuation criteria are not quite in the researcher’s hand. For instance, forward mutations in *rpoB* affect the structure of bacterial RNA polymerase; they are, therefore, non-neutral with regard to the cellular fitness and can affect growth rate to a different extent depending on the genetic background (Figure 1C) [17]. Stress-induced mutagenesis due to the presence of a sub-inhibitory antibiotic concentration during culture growth can increase the death rate, and so can certain genetic backgrounds that severely impact cellular growth (such as mismatch repair deficiency). Mutations in carbon source metabolism-associated genes such as *lacZ* can occur after plating due to adaptive mutagenesis, because the selective agent does not immediately kill the non-mutant cells [18,19]. A phenotypic lag (the phenomenon of delayed expression of a scorable phenotype post mutation due to the necessity to accumulate mutant protein product at sufficient level) is also widespread with mutations causing antibiotic resistance, yet rarely accounted for (Figure 1D) [20,21]. At last, elevated mutation rate can force the researcher to plate only a portion of the culture on selective medium, and because of that, not all mutant cells will be directly detected (imperfect plating, or plating efficiency less than 100%). While in theory in the last example the researcher can stop the culture growth when mutant cells are still countable, this might result in violating the assumption that the final number of cells in culture is much bigger than the starting cell count.

The method of choice for estimating mutation rates by employing the Luria–Delbrück distribution is the Maximum Likelihood Estimation (MLE) [5]. MLE is a statistical tool for finding the value of some parameter of a statistical distribution for which the data are most likely to be observed (drawn from). Based on numerous simulations, Maximum Likelihood-based methods are the most accurate of all available tools for mutation rate estimation under the Luria–Delbrück protocol [22–25]. The popularity of the MLE method can be attributed to Ma, Sandri, and Sarkar, who developed recursive algorithms for efficient computation of the values of the probability mass function (PMF) [26,27].

Among important contributions that allowed to relax some of the strict conditions imposed by the fluctuation protocol that cannot be always easily mitigated by the researcher are the works of: Armitage, who proposed the adjustment for imperfect (not 100%) plating based on binomial thinning (this work was later extended by Crane, Jones, Stewart, Gerrish, Zheng) [9,10,22,28–32]; Angerer, who derived modifications for phenotypic delay and the presence of residual mutations (the subject was also studied by Armitage, Koch, Mandelbrot, Newcombe, Crump et al., and Stewart et al.) [8,10,13,20,33–35]; Mandelbrot, Koch, Jones, and Stewart, who found analytical solutions that allow to account for differential growth of mutant and non-mutant cells (a model that takes into account the relative fitness of the mutant cells compared to non-mutants has been named Mandelbrot–Koch model, see Figure 1C), also with imperfect plating [9,31,33,34]; Zheng, who developed the *B*^0^ distribution, so that the variation between population sizes in parallel sister cultures can be easily corrected for (Figure 1B) [36–39]. Kendall, Zheng, Stewart, Angerer, as well as Dewanji et al. have also considered increased cell death and the effect of the size of the inoculum [8,40–44]. Dewanji et al. proposed an alternative to model mutant birth and death that considers deceleration at the late phase of culture growth. The occurrence of residual mutations is easy to account for if the expected number of post-plating mutants is known, but this in turn is hard to quantify in practice. It has been proposed that adaptive mutagenesis on lactose plates may be limited by the usage of scavenger and/or filler strains and incubating the plates for as short a period of time as possible [19].

### 1.3. Tools for estimating mutation rates

Several tools for mutation rate estimation are currently available: bz-rates [45] and flan [46], both using the so-called Generating Function (GF) method and asymptotical normality assumption, as well as rSalvador [24] and FluCalc [47] based on the Maximum Likelihood (ML) method. bz-rates is an online tool; whereas flan and rSalvador are free packages for an open statistical analysis language R, and FluCalc is written in Python. flan and rSalvador have their online versions available at http://shinyflan.its.manchester.ac.uk and https://websalvador.eeeeeric.com, respectively. A popular tool for estimating mutation rates is FALCOR [48] available at https://lianglab.brocku.ca/FALCOR/index.html, which also uses ML method. Based upon simulations, rSalvador is the most accurate. However, we have found it does not exactly meet our needs. Here, we introduce a novel tool for fluctuation data analysis: <u>MLE</u> <u>Mu</u>tation <u>R</u>ate Calculator, or mlemur.

mlemur is based on the frameworks developed by Qi Zheng. mlemur allows the user to obtain point and interval estimates of mutation rates, compare two datasets using likelihood ratio test to calculate *P* values, and adjust computed *P* values using either Bonferroni, Bonferroni–Holm, or Benjamini–Hochberg corrections, estimate power and sample size for likelihood ratio test, and compute confidence intervals for an arbitrary function of mutation rates.

Unlike flan and rSalvador, mlemur is almost completely typing–free: after initialization via the mlemur::mlemur() command in R, all control is done from within the web browser window. At every stage, mlemur provides the user with information about the type of input required in each data field as well as feedback when an error occurred. Users can also load their own data from an XLS(X) file to compute mutation rates, confidence intervals, *P* values (adjusted or not) for a whole set of strains at once. At last, with a single click, researchers can download the results in a spreadsheet format. Additional options for calculating mutation rates and *P* values for paired data (colony counts on selective and non-selective medium for each culture), finding confidence intervals for an arbitrary function (such as fold or difference) of mutation rates, estimating statistical power (that is, the probability that true differences will be discovered), and determining sample size to achieve a prescribed level of power of the likelihood ratio test, have been implemented.

mlemur can calculate mutation rates using a handful of options:

- the standard Lea–Coulson model (approximate Luria–Delbrück distribution) with almost all the previously mentioned assumptions in place, but overlooking the impact of the inoculum,
- the exact Luria–Delbrück distribution with consideration for the size of inoculum,
- the compound LC–gamma model (*B*^0^ distribution), which accounts for variability in culture sizes,
- the Mandelbrot–Koch model with differential growth of mutants and non-mutants,
- the stochastic modification of Angerer’s model where phenotypic delay is present,
- the Birth–Death (BD) model with cell death,
- the Poisson–LC convolved distribution where post-plating residual mutations are anticipated.

A unique feature of mlemur is that it allows the user to combine most of these generalisations when modelling their data, with either perfect or partial plating. This is especially handy when one scores for multiple phenotypes at once, or when dealing with particularly strong mutators.

Additionally, the functions for simulating fluctuation experiments and calculating mutation rates using a variety of historical methods have been implemented and are available from R console in mlemur. The mathematical background for all developments has been described in detail in Supplemental information, but the most relevant derivations will also be available in the following sections.

In this work we present the possibilities of mlemur and derive new formulas for some extensions of the Luria–Delbrück protocol. We focus on the practical application of the Luria–Delbrück model to estimate mutation rate from simulated and real-world data, investigating the accuracy and confidence interval coverage of point and interval estimates.

## 2. Methods and models

### 2.1. Algorithms for Maximum Likelihood Estimation

The algorithms used in this work for estimating mutation rate are a part of free R statistical package mlemur available at https://github.com/krystianll/mlemur. They are also described in detail in Supplementary File S1.

### 2.2. Simulating fluctuation experiments

Most of the work presented here was done by analysing simulated fluctuation experiments. Simulation algorithm that is a part of mlemur combines previously used approaches with the novel addition of phenotypic lag [38,39,46]. The outline of simulation of a single test tube is presented here:

- Final culture size is taken either as a constant or as a random variable drawn from the gamma distribution.
- Growth (and death if applicable) of the non-mutant cells is assumed to be deterministic and exponential. Time of culture growth is calculated per tube using starting and final number of cells as well as non-mutant death rate. Average number of mutations, which is proportional to the number of cellular divisions, is calculated using average mutation rate, growth rate, death rate, and time of culture growth.
- The actual number of mutations in the test tube is drawn from Poisson distribution using average number of mutations from the previous step.
- Moments of mutation (expressed in terms of number of individual cell divisions in that moment as a fraction of total number of cell divisions at the end of culture growth) are drawn from uniform distribution and then mutation epochs are calculated. For each mutant clone, the length of phenotypic lag is drawn from Poisson distribution with mean being the average length of phenotypic lag (supplied as a number of generations). If a particular mutation epoch exceeds total time of culture growth minus the extent of phenotypic lag, the whole mutant lineage is discarded.
- The size of the mutant clone is drawn from the distribution of the number of cells in a simple birth-and-death process using (8.46) in Bailey 1964 [12].
- If plating is not perfect, the number of mutant colonies on the plates is drawn from binomial distribution.

To simulate the mutant counts under the protein dilution model, a slightly modified version of the code used by Barna was used [49].

The code for simulating a fully stochastic Bartlett model of mutation is an evolution of the Renshaw algorithm described by Zheng [38,44] available at https://github.com/eeeeeric/rSalvador/blob/master/python-examples/simuKessler.py.

### 2.3. Data and code availability

The R and C++ code used for simulating experiments, simulated fluctuation data, and the results of estimations, are available at Zenodo (10.5281/zenodo.7505829). The R package mlemur is available at https://github.com/krystianll/mlemur.

## 3. Results and discussion

### 3.1.

The probability distribution induced by the Luria–Delbrück model

The probability distribution induced by considerations from the Introduction can be written concisely ((18) in [8])

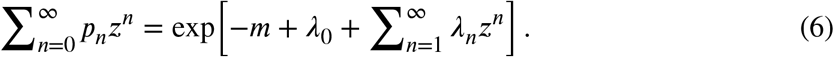

Here *λ_n_* is the expected number of mutations that produce a clone of size *n*, which in case of a simple Luria–Delbrück model is given by (5). It is, however, convenient to re-cast the above as

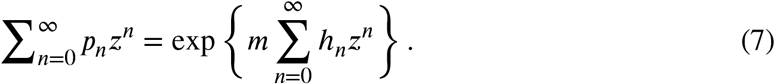

In this form, the auxiliary sequence {*h_n_*} is independent of *m*. Calculating the values of the probability mass function (PMF) comes down to calculating the values of the auxiliary sequence, which needs to be done only once and can be re-used for different values of *m* in each iteration when MLE of *m* is being found numerically by maximisation of the likelihood function.

A formula for {*h_n_*} under the Lea–Coulson model is given by

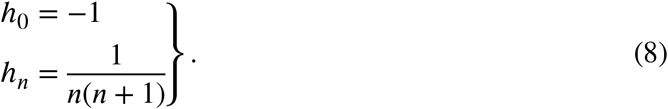

Additionally, when size of the inoculum is to be considered,

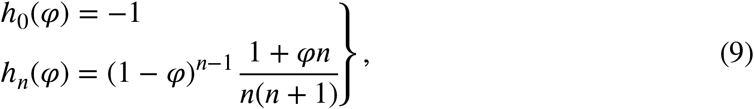

with *φ* = *N_0_/N_t_* (see (7) in [50]). We also note that when substituting *φ* = 0 we arrive at the Lea–Coulson formulation; in other words, Lea–Coulson formulation is a special case of the Luria–Delbrück distribution with *φ* = 0.

In the following sections we will present formulas for {*h_n_*} when some of the assumptions of the Luria–Delbrück model are violated. Many of them were already presented in the literature. We will also present simulation-based studies and, in some cases, real-world examples of how violation of these assumptions affects the mutant distribution and, consequently, estimation of the mutation rates.

### 3.2. Plating a part of the culture

One of the most important deviations from the Luria–Delbrück model concerns plating only a fraction of the culture. A parameter defined as the fraction of culture that was plated is called plating efficiency, *ε*. As argued in [24], mutant cells are not uniformly dispersed throughout the culture, and sampling introduces an additional element of chance. From a statistical point of view, upon sampling each mutant cell undergoes an independent Bernoulli trial with the probability of success (forming a colony) equal *ε*. This is best understood when we consider a culture containing a single mutant cell. If we plate half of this culture, the mutant cell has a 50% chance of being plated. If the culture contains 2 mutant cells, then each of these cells will be plated with a 50% probability: we might expect that one mutant cell will be plated, but it is also possible that we plate 0 or 2, because plating of one cell does not influence the probability of another cell being plated. Armitage was the first to propose to model partial plating using binomial distribution (see (50) in [10]). By multiplying (5) by the expression for PMF of binomial distribution, we can arrive at the formula for expected number of mutations producing *n* mutants, of which *k* are observed (equation (4) in [9]):

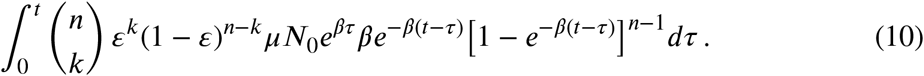

Based on the work of Stewart and Zheng [8,9,22], the formula for {*h_n_*} with imperfect plating assumes the form

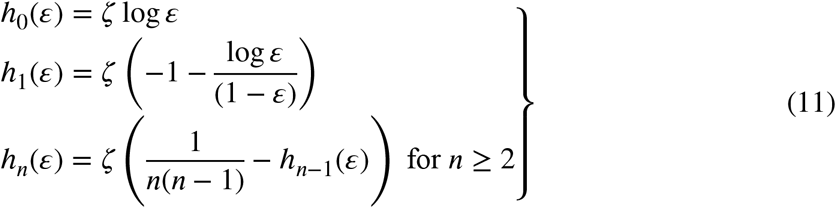

with

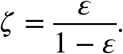

Due to the random nature of sampling, approaches based on dividing either colony counts, or an intermediate estimate *m*\* obtained assuming perfect plating, by the plating efficiency parameter, with the aim to return the true number of mutants in culture or true value of *m*, respectively, will unescapably fail. Figure 2 corroborates the intuitive notion that plating a portion of size *ε* of a culture with the average number of mutations *m*_1_ and plating whole culture with the average number of mutations *m*_2_ = *εm*_1_ give completely different distributions of colony counts. Another popular correction for imperfect plating was proposed by Stewart (equation (41) in [8]):

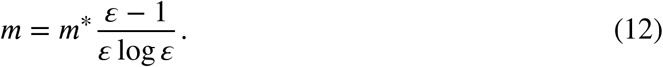

As was shown by Zheng, Stewart’s correction gives acceptable point estimates only for large values of *ε* [24]. However, even then applying it to confidence limits will usually render the more important interval estimates outside nominal coverage (Table 1).

**Figure 2.**
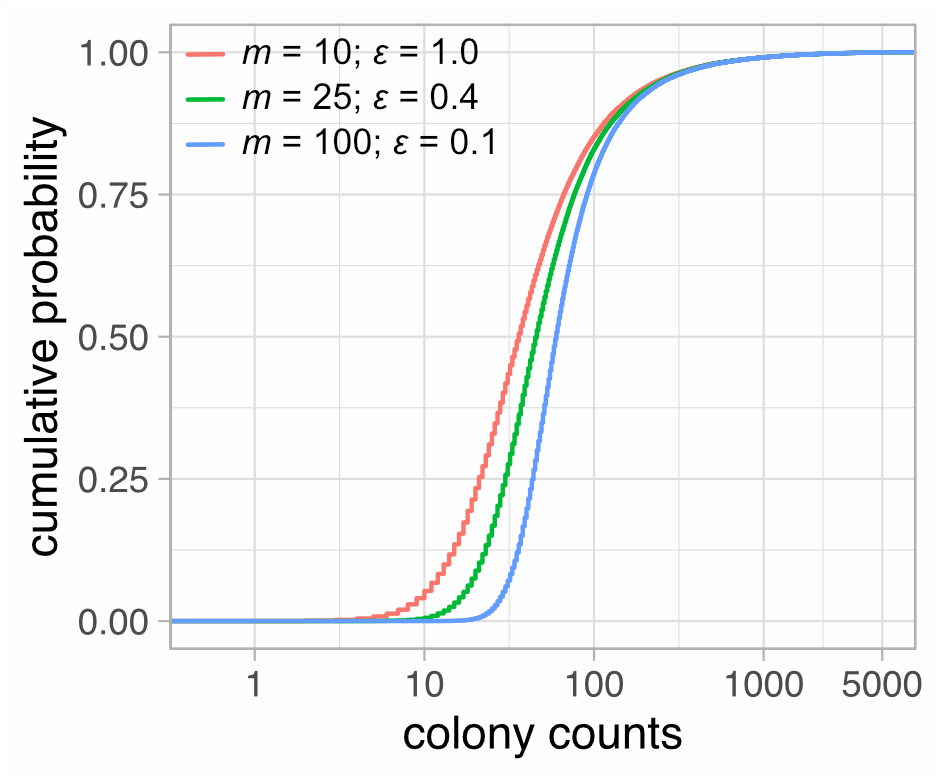
Empirical CDFs of mutant counts under different plating efficiency parameters. One 100,000-tube experiment was simulated for each case. For picture clarity CDFs were censored at the colony count of 5,000. Red — *m* = 10, *ε* = 1; green — *m* = 25, *ε* = 0.4; blue — *m* = 100, *ε* = 0.1.

**Table 1.** Analysis of point and interval estimates for different values of plating efficiency. *N_E_* — number of experiments simulated; *n_C_* – number of cultures in each experiment; *m* — average number of mutations; *ε* – plating efficiency. In all simulations *N*_0_ = 10^3^ and *N_t_* = 10^9^. Nominal CI coverage is 95%. Median CI width was calculated by first expressing the width of CI for each experiment as a percent of corresponding MLE and then taking the median value.

| Simulation parameters |  |  |  | Armitage correction |  |  | Stewart correction |  |  | Rosche and Foster (FALCOR) |  |  |
| --- | --- | --- | --- | --- | --- | --- | --- | --- | --- | --- | --- | --- |
| $N_E$ | $n_C$ | $m$ | $\varepsilon$ | median<br>$m$ | CI<br>coverage | median<br>CI width | median<br>$m$ | CI<br>coverage | median<br>CI width | median<br>$m$ | CI<br>coverage | median<br>CI width |
| 10000 | 30 | 50 | 0.8 | 50.3 | 94.7% | 29.4% | 47.4 | 88.3% | 30.1% | 50.1 | 90.4% | 25.5% |
| 10000 | 30 | 10 | 0.8 | 10.0 | 94.7% | 42.7% | 9.62 | 93.9% | 43.5% | 9.84 | 93.6% | 42.5% |
| 10000 | 30 | 10 | 0.4 | 10.0 | 95.1% | 45.6% | 8.72 | 81.0% | 48.8% | 9.58 | 89.2% | 42.8% |
| 10000 | 30 | 10 | 0.1 | 10.0 | 95.6% | 57.4% | 8.13 | 76.8% | 63.3% | 7.60 | 38.6% | 46.0% |
| 10000 | 30 | 10 | 0.01 | 9.93 | 95.2% | 113% | 8.93 | 95.5% | 118% | 0.600 | 0.00% | 100% |
| 10000 | 100 | 10 | 0.01 | 10.0 | 95.0% | 61.7% | 8.96 | 91.9% | 63.9% | 0.608 | 0.00% | 55.7% |
| 10000 | 30 | 100 | 0.01 | 100 | 94.8% | 43.1% | 66.3 | 4.1% | 54.8% | 70.1 | 9.8% | 23.0% |

Expectedly, sampling from a culture comes with a trade-off: the lower plating efficiency, the wider the confidence interval, and to control the loss of precision of estimation the experimentalist must set up higher number of parallel cultures (Table 1, column: Armitage correction – median CI width).

The popular tool for mutation rate estimation FALCOR cannot properly handle the cases when *ε* < 1. Firstly, the authors propose to normalise the colony counts to 1 mL of the culture, which is invalid for the reasons explained above. In the column “Rosche and Foster” in Table 1, *m* was estimated by first dividing colony counts by *ε* to obtain a theoretical number of mutant cells in the whole culture, and then using these new colony counts for computing MLE with the assumption *ε* = 1. This approach gives acceptable results for bigger values of *ε*, whereas for *ε* = 0.1 *m* is underestimated by ~24% (Table 1, column: Rosche and Foster – median *m*). The underestimation is heavily influenced by the presence of cultures with 0 mutant cells. Secondly, FALCOR utilises the following formula proposed by Rosche and Foster to calculate approximate 95% confidence intervals [5]:

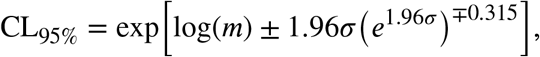

with

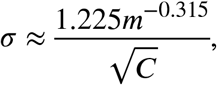

and *C* being the number of cultures. This method works well when the departure from the classical Lea-Coulson model is small, i.e., when the whole, or at least a big part of the culture is plated. Comparison of the coverage of CIs produced using inverted likelihood ratio test with these obtained using Rosche and Foster method shows that the latter are non-conservative especially for very low values of *ε* because the loss of precision caused by partial plating is not accounted for (Table 1, column: Rosche and Foster – CI coverage).

### 3.3. Differential growth of mutants and non-mutants

An important limitation of the classic Luria–Delbrück model is that it requires the mutant cells to have the same growth rate as the wild–type cells, a condition not always satisfied. In particular, when measuring the mutagenesis of the reporter genes that are also essential for the functioning of the cell (such as *rpoB* or *gyrA* in case of *E. coli*), it is expected that the mutations, while obviously favourable during growth on the selective media containing Rif or Nal, might be disadvantageous when cells are not exposed to the selective agent [51].

As far as mutation rates are of concern, accounting for the differential growth rate of mutant and wild-type populations requires only the knowledge about their relative fitness. As a consequence, even a simple strategy of estimating the fitness of two competitors, namely competition assay, will suffice. Other popular fitness assay is, e.g., estimation of the maximum growth rate from OD measurements. For a recent review, the reader may wish to consult [52].

To make things more troublesome, *rpoB* has ~80 mutational sites [53], with different classes of substitutions and sequence contexts; each site may possibly affect cell growth to a different extent. In one *E. coli* study with 8 different *rpoB* mutations it has been shown that the relative fitness can lay anywhere between 0.7 and 1.0 [54]. Similar studies were conducted in other bacteria [55–57]. It is not clear how variability in the mutant fitness affects mutant distribution and accuracy of mutation rate estimation.

The generalisation of the Lea–Coulson model that allows for mutant cells to grow at a different rate that non-mutant cells is often called Mandelbrot–Koch model [33,34,41]. Under this formulation, {*h_n_*} can be written as follows:

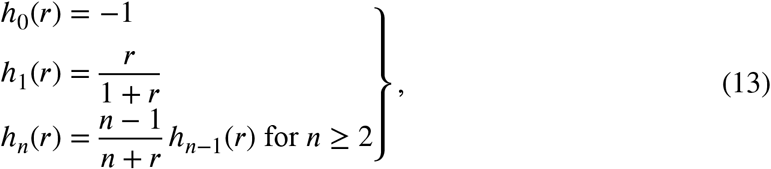

with *r* being the relative growth rate of non-mutants compared to mutants. The reciprocal value *ρ* = 1/*r* is defined as relative growth of mutants to non-mutants. It is easily checked that when substituting *r* = 1 we arrive at the Lea–Coulson formulation.

Solution for the case *ε* ≠ 1 has been found by Stewart and reiterated by Jones [9,31]:

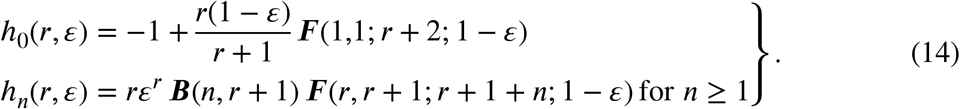

Here, ***B***(*a,b*) is beta function and ***F***(*a,b;c;z*) is Gauss hypergeometric function. To save computational time, {*h_n_*} can be computed using recursive formulas that are provided in the Supplemental Information.

The impact of differential fitness is biggest in case of early mutants which produce more offspring (Figure 1C, Figure 3). For example, lower mutant growth rate results in lower jackpot colony counts and therefore lower variance. As a side note, when *ρ* → 0 the mutant distribution asymptotically becomes Poisson distribution. Table 2 shows that when *ρ* and *ε* are correctly accounted for, both point and interval estimates are accurate for a variety of parameter values.

**Figure 3.**
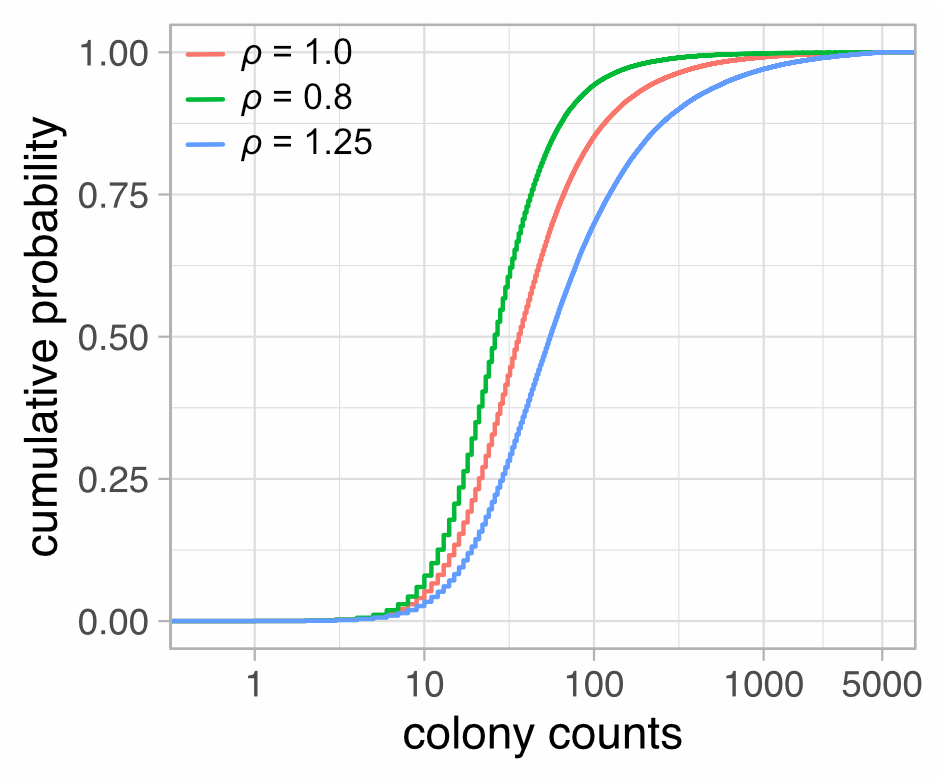
Empirical CDFs of mutant counts with different mutant relative fitnesses. One 100,000-tube experiment with *m* = 10 was simulated for each case. For picture clarity CDFs were censored at the colony count of 5,000. Red — *ρ* = 1.0; green — *ρ* = 0.8; blue — *ρ* = 1.25.

**Table 2.** Analysis of point and interval estimates of *m* for different values of relative mutant fitness and plating efficiency. *N_E_* — number of experiments simulated; *n_C_* — number of cultures in each experiment; *m* — average number of mutations; *ε* — plating efficiency; *ρ* — relative mutant fitness. MK formulation — mutant distribution with correction for *ρ*. In all simulations *N*_0_ = 10^3^ and *N_t_* = 10^9^. LC formulation — simple model where *ρ* = 1. Nominal CI coverage is 95%.

| Simulation parameters |  |  |  |  | MK formulation |  | LC formulation |  |
| --- | --- | --- | --- | --- | --- | --- | --- | --- |
| $N_E$ | $n_C$ | $m$ | $\varepsilon$ | $\rho$ | median<br>$m$ | CI<br>coverage | median<br>$m$ | CI<br>coverage |
| 10000 | 30 | 10 | 0.8 | 1.2 | 10.1 | 95.3% | 12.0 | 65.1% |
| 10000 | 30 | 10 | 0.8 | 0.8 | 10.0 | 95.2% | 8.40 | 62.8% |
| 10000 | 30 | 10 | 0.8 | 0.5 | 10.0 | 95.0% | 6.41 | 0.04% |
| 10000 | 30 | 10 | 0.5 | 1.2 | 10.1 | 94.6% | 12.0 | 65.6% |
| 10000 | 30 | 10 | 0.5 | 0.8 | 10.0 | 94.8% | 8.32 | 62.9% |
| 10000 | 30 | 10 | 0.5 | 0.5 | 10.0 | 94.6% | 6.24 | 0.04% |
| 10000 | 30 | 10 | 0.1 | 1.2 | 10.0 | 95.0% | 12.6 | 64.7% |
| 10000 | 30 | 10 | 0.1 | 0.8 | 10.0 | 94.3% | 7.84 | 63.4% |
| 10000 | 30 | 10 | 0.1 | 0.5 | 9.99 | 95.0% | 5.37 | 0.32% |

### 3.4. Phenotypic lag

Phenotypic lag, or phenotypic delay, is the time required to express the selectable phenotype by cells that already acquired mutation in the marker gene. For example, mutations in *E. coli rpoB* that confer resistance to rifampicin can only be expressed if a sufficient number of wild-type RNA polymerase molecules will be replaced by mutant variants, such that rifampicin does not inhibit transcription to a lethal extent. “Dilution” of the wild-type protein can occur either during cell division, when daughter cell inherits roughly half of the proteins of the parental cell, or due to protein turnover (degradation and resynthesis), although the latter seems to play small role in bacteria. Rif^R^ phenotypic lag of in *E. coli* has been estimated to be 4–5 generations [21]. As a consequence, all mutations occurring in the last 4–5 generations will fail to be expressed (in other words, genotypic mutants will not by phenotypic mutants, Figure 1D). Recently there has been evidence that a big role in phenotypic delay may be played by chromosome segregation in quickly dividing cells [21,58]. Many bacteria in exponential phase start the next cycle of DNA replication before the previous one is finished, leading to a possibility of a heterozygous state where one copy of a gene of interest acquired mutation, whereas another did not. The target gene copy number may vary depending on its location on the chromosome [59]. If selectable allele is recessive, it won’t be expressed until after the cell reaches homozygosity. Simulations performed in the same study suggest that effective polyploidy combined with recessive mutation does not influence mutation rate estimation simply because increased chance of mutation caused by polyploidy cancels out with the phenotypic delay caused by incomplete chromosome segregation [21]. However, effective polyploidy does seem to play a significant role when combined with protein dilution mechanism [60]. Indeed, since the inhibiting activity of antibiotics requires interaction with their molecular targets, it seems that the significance of protein dynamics in the context of phenotypic lag should not be neglected.

The awareness of delayed phenotypic expression of a newly acquired allele is widespread in genetic engineering and seemingly less so when reporting fluctuation data, although its effect on mutant distribution has been considered since the model’s conception. The possible appearance of phenotypic delay was raised in two big fluctuation experiments reported by Newcombe in 1948 and by Boe et al. in 1994 [20,61]. Phenotypic delay was of interest to Armitage, Crump & Hoel, Koch, Angerer, Stewart et al., and Kissling et al. [5,8,10,13,35,58].

The simplest way to model phenotypic lag, which was considered by Armitage and further developed by Angerer, is to assume that a certain amount of time (say, *t*_lag_) since mutational event must pass for the cell to be able to express mutant phenotype [35]. This time can be inputted as a number of generations (with one generation being the time required for population to double in size), because ultimately it is implicitly expressed by the size of the culture at time *t* – *t*_lag_, which we shall denote as *N*_lag_. (We remind here that *t* is the time when culture growth is interrupted, that is, when *N* = *N_t_*; see (1)).

If mutations occurring after *t* – *t*_lag_ are not expressed, it is essentially equivalent to no mutations occurring at all between *t* – *t*_lag_ and *t* (although existing mutants continue proliferating in that time). What this means in practice is that the upper limit of integrals in (5) and (10) becomes *t* – *t*_lag_. The formula derived by Angerer (modified to be applicable to our purposes) is

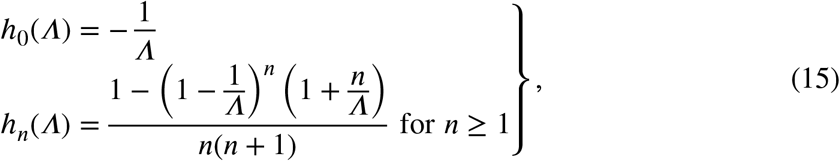

where *Λ* = *N_t_* / *N*_lag_ = *2^l^*. Here *l* denotes the extent of phenotypic lag in generations. Additionally, it is shown in Appendix S1 that when *ε* < 1

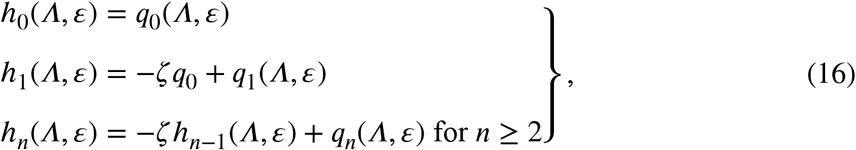

with

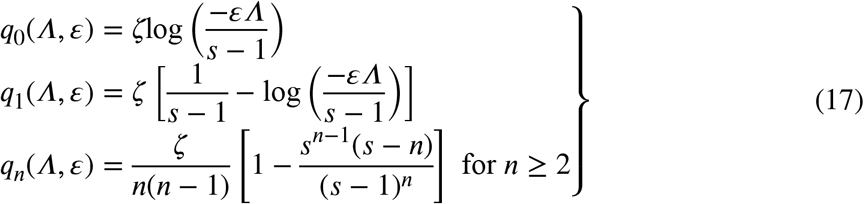

and *s* = *ε* (1 – *Λ*).

What the approach described above does not take into account is the fact that in the Luria–Delbrück model the lifetimes of individual mutant cells are exponentially distributed. From this angle the constant (in terms of time) phenotypic lag seems overly severe: some cell may acquire mutation during lag time but undergo enough divisions such that sensitive protein will be sufficiently diluted. Naturally, the impact will be bigger with longer phenotypic lag and lower mutation rate. To include an element of stochasticity to Angerer model we might assume that the lengths of phenotypic lag *l* of every mutant clone are i.i.d. random variables obeying Poisson distribution with some mean *λ*. The new expressions assume the forms:

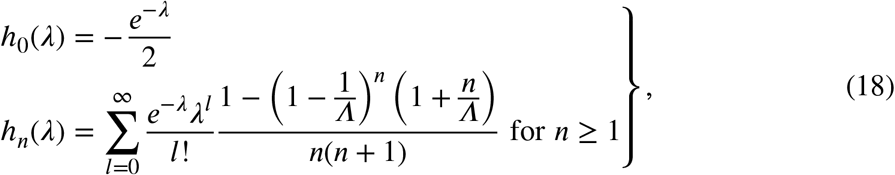

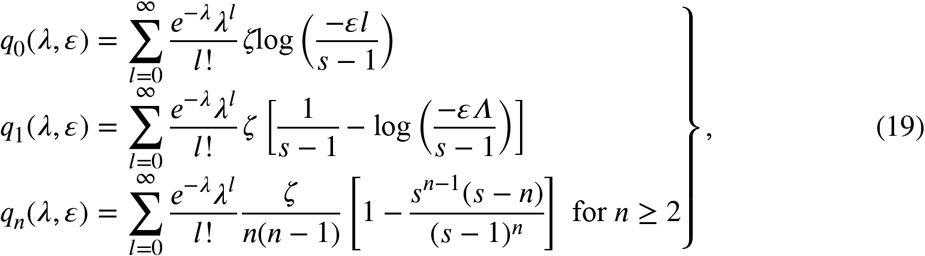

with *Λ* and *s* appropriately substituted.

Delayed phenotypic expression of a mutant phenotype has the biggest impact on the lower tail of mutant distribution [5,10,33], and this fact is reflected in the shapes of empirical CDFs in Figure 4. Accordingly, Table 3 shows that phenotypic lag of just two generations decreases mutation rate estimates by almost a half.

**Figure 4.**
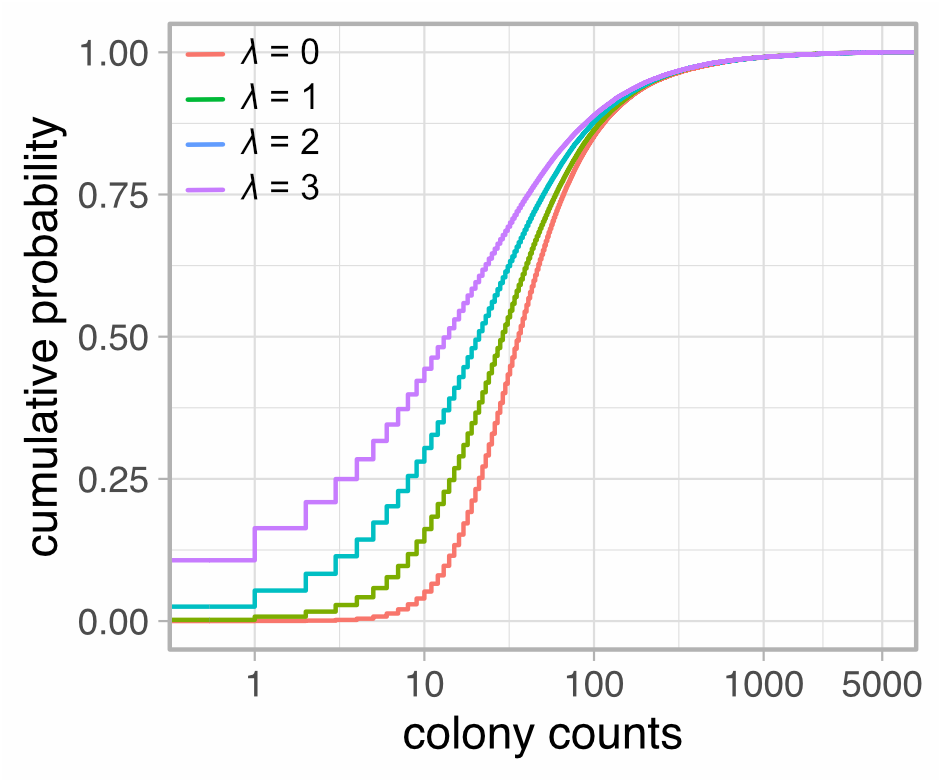
Empirical CDFs of mutant counts with phenotypic lag constant within a clone, but Poisson-distributed from clone to clone. One 100,000-tube experiment with *m* = 10 was simulated for each case. For picture clarity CDFs were censored at the colony count of 5,000. Red — *λ* = 0; green — *λ* = 1; blue — *λ* = 2; purple — *λ* = 3.

**Table 3.** Analysis of point and interval estimates of *m* for different extent of phenotypic lag and plating efficiency. *N_E_* — number of experiments simulated; *n_C_* — number of cultures in each experiment; *m* — average number of mutations; *ε* — plating efficiency; *λ* — average phenotypic lag. In all simulations *N*_0_ = 10^3^ and *N_t_* = 10^9^. SA formulation — stochastic Angerer formulation; mutant distribution with correction for *λ*. LC formulation — simple model where *λ* = 0. Nominal CI coverage is 95%.

| Simulation parameters |  |  |  |  | SA formulation |  | LC formulation |  |
| --- | --- | --- | --- | --- | --- | --- | --- | --- |
| $N_E$ | $n_C$ | $m$ | $\varepsilon$ | $\lambda$ | median $m$ | CI coverage | median $m$ | CI coverage |
| 10000 | 30 | 10 | 1 | 1 | 10.0 | 95.1% | 7.66 | 35.3% |
| 10000 | 30 | 10 | 1 | 2 | 10.0 | 94.8% | 5.41 | 1.32% |
| 10000 | 30 | 10 | 1 | 3 | 10.1 | 94.9% | 3.48 | 0.020% |
| 10000 | 30 | 10 | 0.5 | 1 | 10.0 | 95.2% | 7.80 | 44.7% |
| 10000 | 30 | 10 | 0.5 | 2 | 10.1 | 94.9% | 5.66 | 3.02% |
| 10000 | 30 | 10 | 0.5 | 3 | 10.1 | 95.3% | 3.80 | 0.020% |
| 10000 | 30 | 10 | 0.1 | 1 | 10.0 | 94.7% | 8.16 | 71.6% |
| 10000 | 30 | 10 | 0.1 | 2 | 10.0 | 94.9% | 6.35 | 23.5% |
| 10000 | 30 | 10 | 0.1 | 3 | 9.98 | 94.5% | 4.66 | 2.67% |

To test the performance of the novel stochastic Angerer model against real-world data we revisited two famous experiments performed by Newcombe [20] and by Boe et al. [61]. In the Newcombe study, a total of 8 25-tube (200 μL each) experiments were reported. Each culture was inoculated with either 10^1^ or 10^3^ cells and grown to an average of 3.5×10^8^ cells (see Table 1 in the original paper). Newcombe used two methods to calculate *m* and observed that the *P*_0_ method gave much lower estimate than Luria–Delbrück’s method of the mean. Ruling out the possibility of an upward bias, he suspected that the discrepancy might be a result of phenotypic delay which, again, affects mostly the lower tail of the distribution. Since the *P*_0_ method uses the proportion of cultures containing 0 mutants, it is the most sensitive to bias caused by phenotypic lag.

Several methods of accounting for phenotypic lag have been proposed. One is to simply discard lower tail of the empirical CDF and fit the rest [10]. Another was proposed by Koch: he suggested to divide colony counts by *2^l^* to see the mutant distribution as it was before phenotypic lag started [33]. Newcombe also presented some methods based on the mean number of cells in each culture [20].

Recently, both CDF method and Koch method were employed, using minimisation of the sum of squared errors, to jointly find the value of *m* and the size of the phenotypic lag in Newcombe experiment [49]. The extent of phenotypic delay has been estimated at 3–4 generations, in agreement with analysis by Armitage [10], and the adjusted mutation rate between 1.34×10^−8^ and 2.72×10^−8^.

Here we will employ maximum likelihood estimation under Lea–Coulson and stochastic Angerer models to find the possible values of *m* and *λ*. The MLEs of *μ* under the Lea-Coulson formulation range from 4.27×10^-9^ to 1.13×10^−8^. For all experiments combined, the 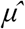 is 6.88×10^-9^ (95% CI, 5.99–7.84×10^-9^). Joint estimation produces *μ* of 7.84×10^-9^ to 5.27×10^−8^ and *λ* between 0.856 and 4.77. When all data are combined, we obtain 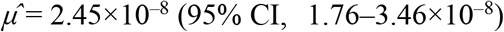 and 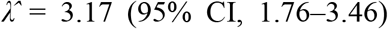. The estimates are therefore in good agreement with previous analyses.

Similar investigation was performed using data reported by Boe et al. [61]. 23 experiments, each comprising 48 parallel tubes, were performed. Cultures were inoculated with a varying number (order of magnitude 10^1^–10^4^) of cells, but this parameter was deemed of little importance. The final size of the culture is not provided in the paper, thus we will focus on the estimates of *m*. Under the Lea–Coulson model 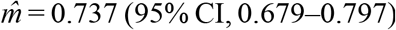, whereas under the stochastic Angerer model it is 0.954 (95% CI, 0.805–1.136), and the estimate of phenotypic lag is 0.601 (95% CI, 0.259–0.957). Similar results were obtained in the Barna analysis (phenotypic lag of 0.6–1) [49].

Barna also performed two experiments with *E. coli* cells growing in defined medium containing either glucose or maltose as a carbon source. This change affects the population doubling time (from ~23 to ~48 min, respectively), but also the estimates of mutation rate under the Lea–Coulson model: 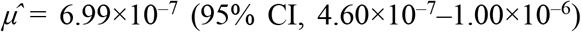 in case of glucose medium and 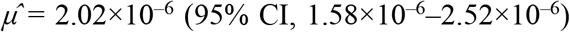 in case of maltose medium. The 3.5-fold difference has been ascribed to the existence of phenotypic lag, which was estimated to be anywhere between 1.58 and 2 generations. Joint estimation of *m* and *λ* shows significant differences only in case of glucose data with the new mutation rate of 1.59×10^-6^ (95% CI, 0.64–4.33×10^-6^) and phenotypic lag of 1.71 (95% CI, 0.03–3.63) generations. Clearly the difference between glucose and maltose data is not significant when one accounts for the phenotypic delay. To conclude, stochastic Angerer model reiterates the results obtained using historical methods.

Considering recent evidence for the significance protein dilution for the expression of a mutant phenotype in bacteria, it is worth investigating how the above model with stochastic phenotypic lag performs against data simulated with consideration for protein dilution. The algorithm for simulations is based on the one from Barna [49] and takes two arguments: the initial number of wild-type proteins *u* in the cell and some threshold value of the number of wild-type proteins in the cell, smaller than *u*, above which the said cell is sensitive to antibiotic. As argued in [60] (see Supplemental Information in the referenced paper), with *u* protein units initially in the cell resistance emerges approximately after *λ* = log_2_*u* generations (see equation S1 in [60]), therefore we have simulated 1, 2, 3, or 5 generations of phenotypic lag by setting *u* to 2, 4, 8, or 32, respectively (Table 4). However, this is only an estimate of when a single randomly chosen cell should develop resistance and does not consider the whole population, in which wild-type protein levels in all cells are co-dependent. If we assume that wild-type proteins need to be completely diluted out of the cell for it to become resistant (by setting threshold value to 0), for every mutant clone, after wild-type proteins are completely distributed into daughter cells, eventually there will be *u* cells possessing single unit of the wild-type protein and thus being sensitive to antibiotic. Therefore, to allow for the whole mutant clone to become resistant, in the simulations the threshold value was set to 1. Since the protein dilution model is indexed by more parameters than the stochastic Angerer model, we considered the possibility that *u* might not easily translate into *λ* and thus *m* and *λ* were estimated jointly. The stochastic Angerer model approximates protein dilution with acceptable precision and accuracy: the error of the point estimation of *m*, as judged by the median, is not bigger than ~5– 10% of the true value, and the confidence intervals retain coverage close to nominal in all cases except when both sample size and phenotypic lag are small (*n_C_* = 30, *u* = 2) (Table 4). This, however, is not caused by the model itself but rather by the well-known inaccuracy of the likelihood ratio test near the boundary of the parameter space: the confidence intervals for *m* cannot be correctly estimated if the value of the nuisance parameter *λ* reaches 0, which is the minimum value *λ* can assume, in agreement with low (~20%) power of the test (Table 4), and this violates the regularity conditions of the Wilks’ theorem [62]. However, the good CI coverage when regularity conditions are met might be related to the fact that the shapes of CDFs under protein dilution model and under stochastic Angerer model with *m* and *λ* chosen to be the same as joint estimates from protein dilution simulated data are remarkably similar (Figure 5).

**Figure 5.**
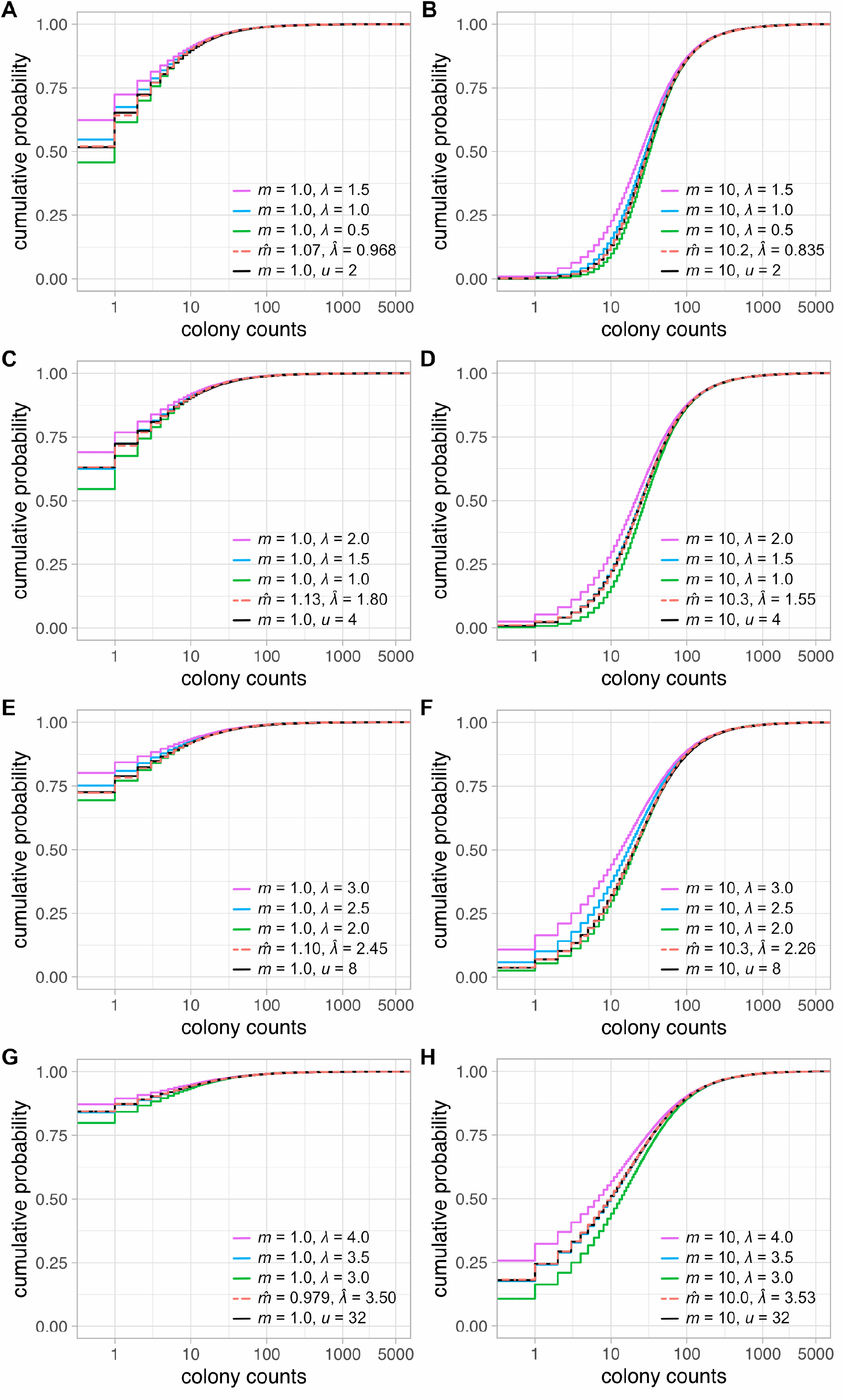
Empirical CDFs of mutant counts under the protein dilution model. One 100,000-tube experiment with *m* = 1 (A, C, E, G) or *m* = 10 (B, D, F, H), and the initial number of protein units cell *u* = 2 (A, B), *u* = 4 (C, D), *u* = 8 (E, F), or *u* = 32 (G, H) that need to be diluted down to 1 or less for the phenotype to be expressed, was simulated for each case, and the empirical CDFs are represented by black continuous lines. MLEs of *m* and *λ* estimated jointly from these data, but assuming stochastic Angerer distribution, were used to simulate additional experiments under stochastic Angerer model, and these CDFs are represented by red dashed lines. Additional CDFs are presented for reference. For picture clarity CDFs were censored at the colony count of 5,000.

**Table 4.** Assessment of efficiency of stochastic Angerer formulation of the Luria–Delbrück distribution against protein dilution model. The threshold value of protein units in the cell above which the phenotype is not expressed has been set to 1. *N_E_* — number of experiments simulated *n_C_* — number of cultures in each experiment; *m* — average number of mutations; *u* — initial number of protein units; *λ* — average phenotypic lag. In all simulations *N*_0_ = 10^3^ and *N_t_* = 10^9^. Joint estimation — MLEs of *m* and *λ* were estimated simultaneously under the stochastic Angerer model. LC formulation —estimation of *m* under the Lea-Coulson formulation. Nominal CI coverage is 95%. Observed power — percent of estimations with *λ* significantly *(p* < 0.05) greater than 0.

| Simulation parameters |  |  |  | Joint estimation |  |  | LC formulation |  | Observed power |
| --- | --- | --- | --- | --- | --- | --- | --- | --- | --- |
| $N_E$ | $n_C$ | $m$ | $u$ | median $m$ | CI coverage | median $\lambda$ | median $m$ | CI coverage | |
| 10000 | 100 | 1 | 2 | 1.06 | 95.1% | 0.938 | 0.703 | 24.6% | 50.6% |
| 10000 | 100 | 1 | 4 | 1.10 | 94.6% | 1.75 | 0.502 | 0.43% | 85.2% |
| 10000 | 100 | 1 | 8 | 1.10 | 95.0% | 2.42 | 0.354 | 0.0% | 93.5% |
| 10000 | 100 | 1 | 32 | 0.97 | 94.9% | 3.46 | 0.179 | 0.0% | 92.8% |
| 10000 | 100 | 10 | 2 | 9.96 | 95.3% | 0.739 | 8.20 | 13.4% | 46.4% |
| 10000 | 100 | 10 | 4 | 10.1 | 94.6% | 1.49 | 6.60 | 0.06% | 91.5% |
| 10000 | 100 | 10 | 8 | 10.3 | 94.9% | 2.22 | 5.07 | 0.0% | 99.8% |
| 10000 | 100 | 10 | 32 | 9.86 | 94.7% | 3.49 | 2.60 | 0.0% | 100% |
| 10000 | 30 | 1 | 2 | 1.06 | 96.8% | 0.900 | 0.699 | 67.0% | 20.9% |
| 10000 | 30 | 1 | 4 | 1.11 | 95.5% | 1.72 | 0.500 | 25.8% | 42.1% |
| 10000 | 30 | 1 | 8 | 1.08 | 95.7% | 2.38 | 0.353 | 5.2% | 52.6% |
| 10000 | 30 | 1 | 32 | 0.95 | 95.6% | 3.40 | 0.183 | 0.0% | 54.3% |
| 10000 | 30 | 10 | 2 | 9.78 | 96.6% | 0.592 | 8.23 | 55.3% | 18.6% |
| 10000 | 30 | 10 | 4 | 9.82 | 95.1% | 1.35 | 6.66 | 11.5% | 47.5% |
| 10000 | 30 | 10 | 8 | 10.0 | 95.1% | 2.12 | 5.11 | 0.78% | 77.3% |
| 10000 | 30 | 10 | 32 | 9.64 | 94.7% | 3.39 | 2.61 | 0.0% | 97.0% |

When the threshold value in the stimulations is set to 0, stochastic Angerer model performs somewhat worse, albeit better than expected, with median *m* being lower than nominal value by about 10% in case of 2–8 protein units and by ~30% in case of 32 protein units, and CI coverage around 90% not only for a more realistic sample size (*n_C_* = 30), but also for a larger one (*n_C_* = 100) which should be more sensitive to deviations from the model assumptions (Table S1). Based on these results as well as good agreement of the estimates with previously published data, stochastic Angerer model may be helpful when analysing data affected by phenotypic delay even though it does not exactly reflect the physiological nature of the process, particularly because the exact distribution of the number resistant bacteria under the protein dilution model (or protein dilution combined with effective polyploidy) is currently unknown.

It should also be noted that there does not seem to be the one correct method to model phenotypic delay: for example, in eukaryotic (particularly mammalian) cells protein degradation plays a bigger role. While arguably some form of accounting for phenotypic delay is better than not correcting at all, it will be interesting to see in the future how a “mechanism-agnostic” formulation such as the one presented above fares against other, more biologically relevant models.

### 3.5. Cell death

Under the classic Luria-Delbruck model the population growth is modelled by a simple pure birth process, i.e., there is no cell death. However, there are certain scenarios where death events can occur frequently, for example:

- Selectable mutation acquired during growth increases the chance of dying of a mutant cell. Wild-type cells are unaffected. This will introduce a negative bias when estimating mutation rate.
- The assayed bacterial strain carries certain genetic mutations that severely affect cellular fitness. Both wild-type and mutant cells will be affected.
- Cells grow in the presence of a sub-inhibitory concentration of an antibiotic. Here again both wild-type and mutant cells can be subjected to dying unless one scores for mutations leading to resistance to the said antibiotic. In such case, only wild-type cells are sensitive, as selective pressure will lead to selection of resistant cells.

If we denote as *β* the growth rate of a strain and by *δ* its death rate, we can also introduce net growth rate of a strain *β*\* = *β* – *δ*. These are assumed constant over time. Further, to ensure population growth we assume that *β* > *δ*. We can also express death rate as a fraction of growth rate, such that *β*\* = *β* – *dβ* = *β*(1 – *d*). Additionally, we can express the death of a cell in terms of probability. If a living cell dies with a probability *p* or divides into two daughter cells with a probability 1 – *p*, then *β*\* = (1 – *p*)*β* – *dpβ*. From this relation it follows that the relative death rate *d* is the same as the odds of dying of a cell: *d* = *p*/(1 – *p*), and the net growth rate can be expressed as *β*\* = *β*(1 – 2*p*)/(1 – *p*). Since we assumed *d* < 1, clearly *p* < 0.5.

We can extend our assumption of deterministic growth of non–mutant cells by saying that these cells *grow and die* deterministically, which simply means that *β* in (1) and (3) is replaced by *β*\*. This should hold for reasonable values of *d* and sufficiently big inocula. For example, under a simple stochastic model, the chance of extinction as *t*→∞ of a population starting with *N*_0_ cells is *d*^*N*0^ (see (8.59) in [12]), so for example for *d* = 0.9 and *N*_0_ = 1000 the chance of extinction is ~10^-46^. How stochastic growth of non-mutant cells affects the estimates of mutation rates will be discussed in the next section.

It is important to remind that in our model cells mutate during division (hence the name mutation rate per cell division) and thus expressions (1) and (3) are of use only when the number of divisions can be equated to *N_t_* – *N*_0_. This is obviously not the case when there is cell death as in this scenario some divisions are masked by dying cells. Simply put, with cell death more divisions are required to reach the same number of cells (Figure 1F). The real number of cell divisions can be extracted by simple reasoning. If the chance of death of a wild-type cell is *p*, out of a given number of wild-type cell events a fraction *p* will be deaths and 1 – *p* will be births; for example when *p* = 0.4 (which implies *d* = 2/3), out of 10 events we will have 6 births and 4 deaths, so starting from a single (*N*_0_ = 1) cell we have *N_t_* = 1 +1 +1 +1 +1 +1 +1 −1 −1 −1 −1 = 3. The increase in cell count is *N_t_* – *N*_0_ = 2. On the other hand, *N_t_* – *N*_0_ = Births – Deaths, and Births/Deaths = (1 – *p*)/*p* = 1/*d*, so the number of cell divisions is clearly Births = (*N_t_* – *N*_0_)/(1 – *d*) = 6. Hence, when there is wild-type cell death, mutation rates are inflated by a constant factor (1 – *d*)^-1^. The correcting factor was first derived by Newcombe [20] and reiterated by Zheng [44]. The phenomenon of overestimation of mutation rates due to cell death was also reported by Frenoy & Bonhoeffer based on similar reasoning supported by simulations [63].

The dynamics of the mutant cells can be modelled by a simple birth-and-death process, as previously suggested [34,41,42,64,65]. Under this model, upon completion of the lifetime, a cell can either divide (+1) or die (−1). The most profound difference between mutant and non-mutant cells is that each mutant clone starts with a single cell, therefore its chance of dying cannot be neglected: if the original mutant cell dies, the whole mutant lineage will never be. As a consequence, mutant cell death introduces a downward bias on mutation rate estimation (Figure 1E).

For reasons explained later we will focus on the situation where death rate is the same for wild-type and mutant cells, that is, *β_1_*/β_2_\** = *β_1_/β_2_* = *r*. (The case when *d_1_* ≠ *d_2_*, for example when cells are exposed to a sub-inhibitory concentration of an antibiotic and *d*_2_ = 0, was studied by Zheng [44]). The expression for the auxiliary sequence {*h_n_*} assumes the form

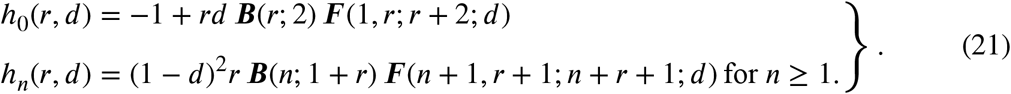

When only a part of the culture is plated, the expression is somewhat more complicated:

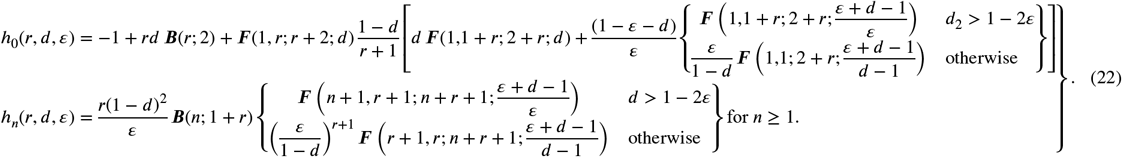

Simulations show that the upward bias from increased number of divisions has a dominant effect over the downward bias caused by dying mutant clones, resulting in higher mutant counts (Figure 6) as well as overestimated mutation rates (Table 5). Small death rates (< 10% of growth rate) have little impact on the point estimates of mutation rates. However, the bias quickly begins to rise and at *d* = 0.25 the difference is ~20%. At *d* = 0.6 – 0.7 mutation rates are essentially doubled (Table 5).

**Figure 6.**
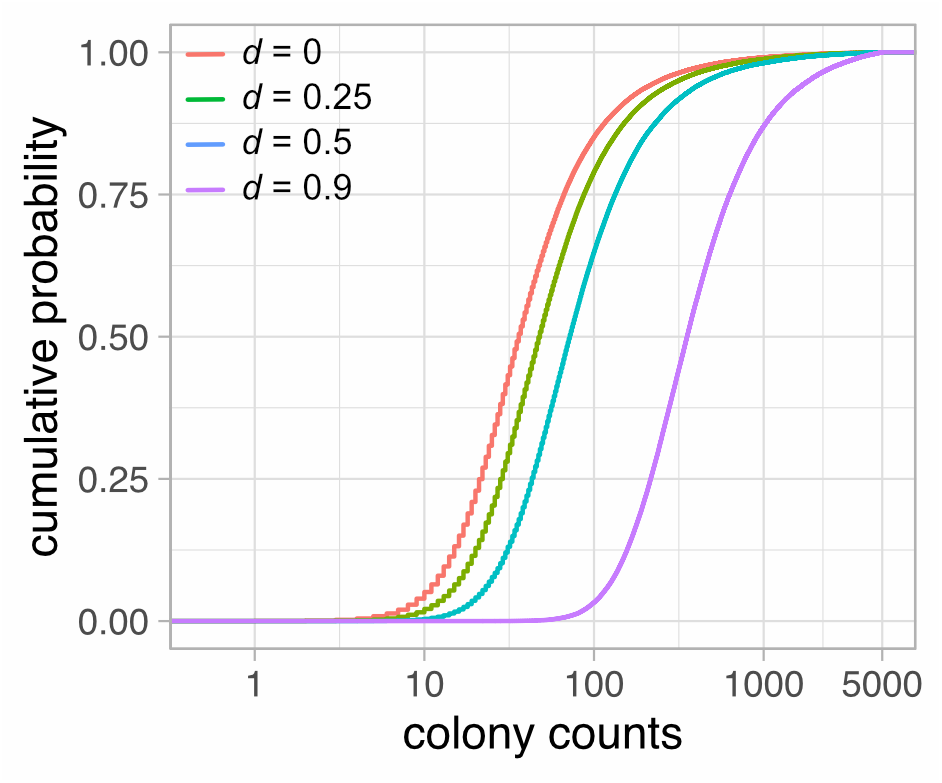
Empirical CDFs of mutant counts with non-zero death rate. One 100,000-tube experiment with *m* = 10 was simulated for each case. The death rate is the same for wild-type and mutant cells. For picture clarity CDFs were censored at the colony count of 5,000. Red — *d* = 0; green — *d* = 0.25; blue — *d* = 0.5; purple — *d* = 0.9.4.

**Table 5.** Analysis of point and interval estimates of *m* under conditions of a non-zero death rate. *N_E_* — number of experiments simulated; *n_C_* — number of cultures in each experiment; *m* — average number of mutations; *ε* — plating efficiency; *d* — relative death rate. In all simulations *N*_0_ = 10^3^ and *N_t_* = 10^9^. BD formulation — Luria–Delbrück model with mutant dynamics modelled using simple birth-death process. LC formulation — simple model where *d* = 0. Nominal CI coverage is 95%.

| Simulation parameters |  |  |  |  | BD formulation |  | LC formulation |  |
| --- | --- | --- | --- | --- | --- | --- | --- | --- |
| $N_E$ | $n_C$ | $m$ | $\varepsilon$ | $d$ | median $m$ | CI coverage | median $m$ | CI coverage |
| 10000 | 30 | 10 | 1 | 0.1 | 10.0 | 95.0% | 10.8 | 89.5% |
| 10000 | 30 | 10 | 1 | 0.25 | 10.0 | 94.6% | 12.3 | 53.8% |
| 10000 | 30 | 10 | 1 | 0.65 | 10.1 | 94.9% | 21.2 | 0.0% |
| 10000 | 30 | 10 | 1 | 0.9 | 10.0 | 94.8% | 52.7 | 0.0% |
| 10000 | 30 | 1 | 1 | 0.1 | 1.00 | 94.8% | 1.06 | 93.5% |
| 10000 | 30 | 1 | 1 | 0.25 | 1.00 | 95.1% | 1.18 | 87.2% |
| 10000 | 30 | 1 | 1 | 0.65 | 1.00 | 94.5% | 1.79 | 19.5% |
| 10000 | 30 | 1 | 1 | 0.9 | 1.00 | 94.8% | 3.52 | 0.0% |
| 10000 | 30 | 10 | 0.1 | 0.1 | 10.0 | 94.8% | 10.8 | 91.9% |
| 10000 | 30 | 10 | 0.1 | 0.25 | 10.0 | 94.7% | 12.4 | 67.2% |
| 10000 | 30 | 10 | 0.1 | 0.65 | 10.1 | 94.6% | 22.0 | 0.0% |
| 10000 | 30 | 10 | 0.1 | 0.9 | 10.1 | 95.0% | 56.0 | 0.0% |

At the end we would like to discuss the case when the presence of an antibiotic in the liquid medium promotes selection of resistant bacteria (*d*_1_ ≠ *d*_2_ = 0), which was studied in [44]. Under the assumptions considered above, bacterial growth in the presence of a sub-inhibitory concentration of an antibiotic has, apart from increasing the number of cell divisions, the additional effect of decreasing the net growth rate of the non-mutant cells, leading to *ρ*\* > 1 and thus further inflating mutant counts. (This scenario was not discussed by Frenoy & Bonhoeffer as in their algorithm mutants grow with rate *β*(1 – *d*) [63]). However, setting very big wild-type death rate such as *d_1_* = 0.95 leads to *ρ*\* = 20. When mutant population grows so much faster than the wild-type cells, one can easily encounter a situation where mutant count is much bigger than that of non-mutants, violating the requirement that mutant cells comprise only a negligible part of the culture. Additionally, the switch to phenotypical resistance is probably not instantaneous, and in reality, mutant cells will continue to die with the same rate as wild-type cells for the period of phenotypic lag, or possibly with gradual change of death rate from that of wild-type cells to that of mutants (in any case, death rate is probably not constant over time). As departing from these useful assumptions would inevitably lead to a significantly more complicated model, for the time being mlemur cannot treat such cases.

### 3.6. Variation of the final number of cells in parallel cultures

Under the Luria–Delbrück model, we assume that each sister culture contains the same number of cells. It is clear, however, that complete homogeneity of the culture sizes is impossible to achieve. This problem has been studied previously by Ycart and Veziris as well as Zheng [38,39,66]. In essence, because the mutation rate is constant, smaller number of cells in a given culture will result in smaller average number of mutations (see (2)), which in turn will increase the fraction of smaller colony counts. As the values of the PMF of the Luria–Delbrück distribution are, in general, bigger for smaller colony counts, we can deduce that high variability in culture size will deflate the estimate of mutation rate.

A good unitless measure of dispersion is coefficient of variation *(CV)*, which is standard deviation (SD) divided by mean number of cells in each culture (*N_t_*). Two studies were conducted to assess the impact of *CV* on the estimates of *m*. In the study by Ycart and Veziris, the fluctuation data were simulated assuming a log-normal, gamma, or other similarly shaped distribution [39]. In Zheng 2016 each culture was simulated using a Bartlett stochastic algorithm [38]. The conclusion from these works was that *CV* as big as 0.2 has an insignificant impact on the estimates of *m*.

When *CV* is big, Zheng proposed two methods to deal with extra variability. One is to use *B*^0^ distribution, which is a mixture of gamma and Luria–Delbrück distributions indexed by *m* and *CV*[37,38]. While mutant counts are Luria–Delbrück distributed, the parameter *m* is itself a random variable obeying the gamma distribution, and *CV* of *m* is assumed to be the same as *CV* of *N_t_*. Another method is to use the so-called “Golden Benchmark method” which simply means estimating *μ* directly using pairs of mutant counts and population sizes for each test tube [38].

In the current study, we investigated the impact of *CV* in the context of wild-type cell death. The model developed in the previous section neglects the stochastic nature of the wild-type dynamics, whose important consequence is inflation of the variance of culture size. If we assume that wild-types grow according to the simple birth and death process, then using (8.48) and (8.49) in [12] and with small rearrangements we arrive at the following formula for the coefficient of variation of *N_t_*:

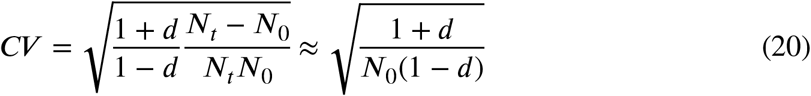

which shows that the coefficient of variation increases with smaller *N*_0_ and bigger *d*. However, setting for example *N*_0_ = 10^3^, *N_t_* = 10^8^, and *d* = 0.7 gives *CV* = 0.075. Even *d* as big as 0.95 gives *CV* = 0.197, within the 20% safety threshold [38], indicating that the variability might be controlled so long as inoculum is sufficiently big.

To test whether one can model the dynamics of the wild-types using a deterministic approach with acceptable results, we simulated fluctuation data using the fully stochastic fixed-time Bartlett algorithm by Qi Zheng which can be found in the rSalvador source files and was described in [44]. The fate of the culture is tracked event-by-event, and each event might be one of four: wild-type birth, wild-type death, mutant birth, or mutant death, with mutant birth resulting from either cell division or mutation. The lifetimes of all cells are exponentially distributed. Consequently, the number of births and deaths, as well as their ratio and *N_t_*, are now all random variables (because the process is stochastic). However, with sufficiently long growth time the mean number of births should be close to its deterministic equivalent, which, as we have shown previously, is given by (*N_t_* – *N*_0_)/(1 – *d)*. For example, if we set *N*_0_ = 100, *t* = 15.4, *μ* = 10^-4^, and *d* = 0.7 (which corresponds to *β*_1_ = 1, *β*_2_ = 1, *δ*_1_ = 0.7, *δ*_2_ = 0.7 in the simulation algorithm), the average *N_t_* from 10^6^ simulated test tubes is 10,086, and the mean number of births is 33,287. The same number of births is produced in the deterministic setting as calculated using (10,086 – 100)/(1 – 0.7).

We have simulated a series of 10,000 100-tube experiments with the values of *d* between 0.5 and 0.9, *μ* = 10^-4^ or 10^-5^, and *t* chosen such that *N_t_* varied from 10^4^ to 2.5’10^5^ (which corresponded to *m* between 1 and 25). Since *N_t_* was kept relatively small to save computational time, and in some cases *φ* was as big as 0.02, we choose to correct for the size of inoculum in our estimations of *m*. However, in a real-world scenario one does not have access to the number of wild-type cells, because the number of colonies on non-selective plates is a sum of mutant (*n*) and non-mutant colonies (*N_t_*). Therefore, *μ* was estimated using average total culture size: to obtain the estimate, we divided *m* by (*N_t_* + *n* – *N*_0_)/(1 – *d*).

Our experiments demonstrate that when the number of wild-types is a random variable (because of their stochastic growth), *μ* is slightly underestimated when one uses the Luria–Delbrück distribution with cell death, as judged by median *μ* lower by 1–7% than the nominal value. The results show that the performance of both point and interval estimates strongly depends on the size of inoculum, as this parameter greatly affects *CV*, but also on *N_t_*, or rather on the associated *m* (Table 6, column LD distribution). For example, for *d* = 0.7, *μ* = 10^-4^, and *N*_0_ = 100, the 95% CI coverage when *N_t_* ≈ 10^4^ (*m* ≈ 1) is 94.9%, but when *N_t_* ≈ 5×10^4^ (*m* ≈ 5) it is 86.7% and for *N_t_* ≈ 10^5^ (*m* = 10) it drops to 77.4%, even though in all these cases *CV* remains stably around 24% (Table 6, lines 5–8). The likely explanation for this observation is that bigger values of *m* produce higher per-plate mutant counts, increasing precision, and this in turn negatively affects CI coverage; for example, decreasing mutation rate to 10^−5^ while keeping *N_t_* ≈ 10^5^ (*m* ≈ 1) increases CI coverage to 94.7% (Table 6, compare lines 8 & 9). The drop in accuracy might be alleviated by decreasing the variability in culture sizes: for *μ* = 10^−4^ and *N_t_* ≈ 10^5^ (*m* ≈ 10), increasing *N*_0_ to 250 (*CV* ≈ 15%) gives 95% CI coverage 92.0%, and for *N*_0_ = 500 (*CV* ≈ 11%) it is 93.8% (Table 6, compare line 8 to lines 11–12). Similar observations can be made for the case where *N_t_* ≈ 5×10^4^ (*m* ≈ 5) (Table 6, compare lines 6 & 10).

**Table 6.** Point and interval estimates of *μ* under a fully stochastic model with cell death. *N_E_* — number of experiments simulated; *n_C_* — number of cultures in each experiment; *N*_0_ — size of inoculum; *t* — time of culture growth; *μ* — mutation rate per cell; *d* — relative death rate; *ρ* — relative mutant fitness; *N_t_* + *n* — average final number of cells in the culture (non-mutant and mutant). Cell cultures were simulated using a fully stochastic Bartlett algorithm until a prescribed value of *N_t_* + *n* was reached but modelled using a simple birth-death process for mutants with deterministic growth of wild-types, with consideration for the inoculum, using either Luria–Delbrück distribution or *B*^0^ distribution where *CV* is taken into account. For each experiment, *μ* was calculated by dividing the estimate of *m* by (*N_t_* + *n* – *N*_0_)/(1 – *d)*. Mean and *CV* for *N_t_* was calculated for each experiment, and median values are presented in the table. Nominal CI coverage is 95%.

| Line | Simulation parameters | | | | | | | Culture size | | LD distribution | | $B^0$ distribution | | Golden Benchmark | |
| --- | --- | --- | --- | --- | --- | --- | --- | --- | --- | --- | --- | --- | --- | --- | --- |
| | $N_E$ | $n_C$ | $N_0$ | $t$ | $\mu$ | $d$ | $\rho$ | median<br>$N_t + n$ | median<br>$CV$ | median $\mu$ | CI<br>coverage | median $\mu$ | CI<br>coverage | median $\mu$ | CI<br>coverage |
| 1 | 10000 | 100 | 100 | 9.2 | $1 \times 10^{-4}$ | 0.5 | 1 | $9.96 \times 10^3$ | 17.2% | $9.87 \times 10^{-5}$ | 95.4% | $1.00 \times 10^{-4}$ | 95.6% | $9.99 \times 10^{-5}$ | 95.3% |
| 2 | 10000 | 100 | 100 | 12.4 | $1 \times 10^{-4}$ | 0.5 | 1 | $4.93 \times 10^4$ | 17.2% | $9.76 \times 10^{-5}$ | 93.3% | $1.00 \times 10^{-4}$ | 95.7% | $1.00 \times 10^{-4}$ | 95.1% |
| 3 | 10000 | 100 | 100 | 13.8 | $1 \times 10^{-4}$ | 0.5 | 1 | $9.94 \times 10^4$ | 17.2% | $9.69 \times 10^{-5}$ | 90.4% | $1.00 \times 10^{-4}$ | 95.9% | $1.00 \times 10^{-4}$ | 94.9% |
| 4 | 10000 | 100 | 200 | 12.4 | $1 \times 10^{-4}$ | 0.5 | 1 | $9.87 \times 10^4$ | 12.2% | $9.85 \times 10^{-5}$ | 93.6% | $1.00 \times 10^{-4}$ | 95.7% | $1.00 \times 10^{-4}$ | 95.1% |
| 5 | 10000 | 100 | 100 | 15.4 | $1 \times 10^{-4}$ | 0.7 | 1 | $1.02 \times 10^4$ | 23.6% | $9.75 \times 10^{-5}$ | 94.9% | $1.00 \times 10^{-4}$ | 96.1% | $1.00 \times 10^{-4}$ | 95.6% |
| 6 | 10000 | 100 | 100 | 20.7 | $1 \times 10^{-4}$ | 0.7 | 1 | $4.99 \times 10^4$ | 23.7% | $9.50 \times 10^{-5}$ | 86.7% | $1.00 \times 10^{-4}$ | 96.0% | $1.00 \times 10^{-4}$ | 94.9% |
| 7 | 10000 | 100 | 100 | 20.7 | $1 \times 10^{-4}$ | 0.7 | 0.8 | $4.98 \times 10^4$ | 23.7% | $9.53 \times 10^{-5}$ | 86.4% | $1.00 \times 10^{-4}$ | 96.2% | $1.00 \times 10^{-4}$ | 94.8% |
| 8 | 10000 | 100 | 100 | 23 | $1 \times 10^{-4}$ | 0.7 | 1 | $9.95 \times 10^4$ | 23.7% | $9.38 \times 10^{-5}$ | 77.4% | $9.99 \times 10^{-5}$ | 96.4% | $1.00 \times 10^{-4}$ | 94.9% |
| 9 | 10000 | 100 | 100 | 23 | $1 \times 10^{-5}$ | 0.7 | 1 | $9.92 \times 10^4$ | 23.7% | $9.71 \times 10^{-6}$ | 94.7% | $9.98 \times 10^{-6}$ | 95.9% | $9.99 \times 10^{-6}$ | 95.2% |
| 10 | 10000 | 100 | 250 | 17.7 | $1 \times 10^{-4}$ | 0.7 | 1 | $5.07 \times 10^4$ | 15.0% | $9.81 \times 10^{-5}$ | 93.5% | $1.00 \times 10^{-4}$ | 95.6% | $1.00 \times 10^{-4}$ | 94.8% |
| 11 | 10000 | 100 | 250 | 20 | $1 \times 10^{-4}$ | 0.7 | 1 | $1.01 \times 10^5$ | 15.0% | $9.75 \times 10^{-5}$ | 92.0% | $1.00 \times 10^{-4}$ | 95.5% | $1.00 \times 10^{-4}$ | 95.0% |
| 12 | 10000 | 100 | 500 | 17.7 | $1 \times 10^{-4}$ | 0.7 | 1 | $1.01 \times 10^5$ | 10.6% | $9.88 \times 10^{-5}$ | 93.8% | $1.00 \times 10^{-4}$ | 95.2% | $1.00 \times 10^{-4}$ | 95.0% |
| 13 | 10000 | 100 | 250 | 53 | $1 \times 10^{-4}$ | 0.9 | 1 | $5.04 \times 10^4$ | 27.3% | $9.26 \times 10^{-5}$ | 75.7% | $1.00 \times 10^{-4}$ | 96.6% | $1.00 \times 10^{-4}$ | 94.8% |
| 14 | 10000 | 100 | 500 | 46.1 | $1 \times 10^{-4}$ | 0.9 | 1 | $5.05 \times 10^4$ | 19.3% | $9.64 \times 10^{-5}$ | 89.9% | $1.00 \times 10^{-4}$ | 95.9% | $9.99 \times 10^{-5}$ | 95.0% |
| 15 | 10000 | 100 | 500 | 53 | $1 \times 10^{-4}$ | 0.9 | 1 | $1.01 \times 10^5$ | 19.4% | $9.56 \times 10^{-5}$ | 85.1% | $9.99 \times 10^{-5}$ | 96.1% | $9.99 \times 10^{-5}$ | 95.4% |
| 16 | 10000 | 100 | 500 | 53 | $1 \times 10^{-5}$ | 0.9 | 1 | $1.00 \times 10^5$ | 19.4% | $9.78 \times 10^{-6}$ | 94.5% | $1.00 \times 10^{-5}$ | 95.4% | $1.00 \times 10^{-5}$ | 95.0% |
| 17 | 10000 | 100 | 1000 | 46.1 | $1 \times 10^{-4}$ | 0.9 | 1 | $1.01 \times 10^5$ | 13.7% | $9.79 \times 10^{-5}$ | 92.3% | $1.00 \times 10^{-4}$ | 95.5% | $9.99 \times 10^{-5}$ | 94.7% |
| 18 | 10000 | 100 | 1000 | 55.2 | $1 \times 10^{-4}$ | 0.9 | 1 | $2.51 \times 10^5$ | 13.7% | $9.74 \times 10^{-5}$ | 89.2% | $9.99 \times 10^{-5}$ | 96.0% | $9.98 \times 10^{-5}$ | 95.1% |
| 19 | 10000 | 100 | 2000 | 39.1 | $1 \times 10^{-4}$ | 0.9 | 1 | $1.00 \times 10^5$ | 9.6% | $9.89 \times 10^{-5}$ | 94.5% | $9.99 \times 10^{-5}$ | 95.7% | $9.99 \times 10^{-5}$ | 95.4% |

The usage of *B*^0^ distribution significantly improves the accuracy of point estimates of *μ* at any value of *CV*, albeit at the cost of producing too conservative confidence intervals when *CV* and/or *m* is high (we observe 95% CI coverage at 95.2–96.6%, Table 6, column *B*^0^ distribution). This problem only affects the CIs of *μ*, and the confidence intervals of *m* retain the correct coverage; for example, if in the estimations described in Line 8 of Table 6 we replace each observed *N_t_* with the mean from 10^6^ test tubes (pooled from all experiments) equal 9.95×10^4^, the CI coverage becomes 95.0%. The cause of this discrepancy is that it is the value of *m* that is directly estimated by the algorithm, and the variability in *N_t_* is reflected is a similar variability in *m*, which is already incorporated in the model. Dividing *m* by the true value of *N_t_* results in a situation such that we correct for the variability in culture sizes twice. Consequently, values of *m* farther from the true value are partially offset by similar variability of *N_t_*, and thus the actual observed variance of *μ* is smaller than anticipated and the CI coverage increases.

Alternatively, if the values of *N_t_* for each culture are known, one can also use the Golden Benchmark method described in previous paragraphs: we note a high accuracy of point estimates as well as CI coverage at 94.7%–95.6% (Table 6, column Golden Benchmark). Good performance of the Golden Benchmark method is particularly striking in light of the fact that we disregarded the variability in the proportion of deaths *vs*. births, replacing it with its mean value equal *d*. This parameter plays a crucial role in recovering the number of cell divisions from the number of living cells. However, a brief analysis of the number of births and deaths in the simulated data shows that the distribution of the deaths-to-births ratio is characterised by a low variance, which might explain why this parameter seems to have a negligible impact on the estimates in practice (Table S2, Figure S1).

Our observations remain consistent when *d* = 0.5 and also when *d* = 0.9, although in the latter case one needs to use bigger inocula (order of magnitude 10^3^) to counterbalance the extra variability caused by high death rate and its effect on the estimates using the Luria–Delbrück distribution (Table 6). Decreasing the sample from 100 to a more realistic 30 cultures per experiment improved the coverage of confidence intervals produced using Luria–Delbrück distribution, which is most evident when one compares, for example, Lines 6, 7, 8, 13, and 15 between Table 6 and Table S3. Overall, these results suggest that it is possible to neglect the stochastic nature of the wild-type cell dynamics without much loss of accuracy and precision of the estimation so long as the values of *m* and *CV* are controlled, and cultures are grown for a sufficiently long time.

### 3.7. A universal sequence?

Feeling encouraged by the number of developments concerning relaxation of the requirements of the Luria–Delbrück protocol, it might be appealing to attempt derivation of a universal sequence that combines existing generalisations of the Lea–Coulson model. However, one quickly arrives at complex dilemmas, one of which has been described in section 3.6. A related problem can be easily imagined if a researcher wishes to correct simultaneously for differential growth of mutant and wild-type cells as well as phenotypic lag. If for a period of time a genetic mutant does not express resistant phenotype, we can imagine that the same trait will be extended to other phenotypes: growth rate and death rate. In the most naïve scenario, a living cell will instantly change its growth dynamics from that of a wild-type cell to that of a mutant when the period of phenotypic lag expires. A closed-form expression for the case *d* = 0 can be derived under this assumption, because the marginal distribution of the number of cells starting from 1 and growing with a suddenly switching growth rate is simply

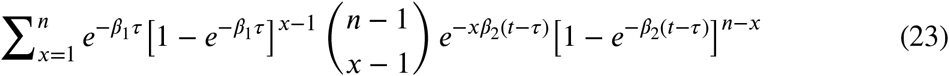

with *τ* being the length of phenotypic lag. The expression becomes significantly more complicated when we replace the Yule process in the above with the simple birth-and-death process. In any case, it seems more realistic that the change in both growth rate and death rate would be gradual over the period of phenotypic delay as wild-type proteins are being successively replaced by their mutant forms. We could therefore, instead of taking *β* and *δ* as constants, model mutant dynamics using a non-homogeneous process such as the one described in chapter 9.3 in Bailey [12]. This however raises a question of how exactly the change in *β* and *δ* over time occurs.

Clearly the outcome of a fluctuation assay may be affected by more than one element at the same time and thus it is desirable for the statistical model to include as many of these elements as possible. Alas, the progress in this area is limited by lack of biological data as well as increasing complexity of the model and associated with it mathematical and computational difficulties. Hence, the adjustment for phenotypic lag is currently limited to cases where we assume that apart from acquiring the ability to grow on selective medium there are no other changes in phenotype (i.e., *β*_1_ = *β*_2_ and *δ*_1_ = *δ*_2_).

### 3.8. Concluding remarks

80 years after publication of the Luria & Delbrück paper, fluctuation analysis has seen numerous developments that aim to relax its strict requirements. For a long time, however, the researchers were limited to a very basic protocol that did not account for inter-strain differences in fitness, phenotypic lag, cellular death rate, and other deviations that affect the number of mutant colonies on the plate. One of such deviations is imperfect plating, which should be modelled properly to obtain correct results (Table 1). To this day the nuances are frequently disregarded, perhaps due to a lack of knowledge, judging by the unfailing popularity of FALCOR (e.g., [67–69]). With the computational power of modern CPUs, it is becoming feasible to incorporate multiple additional parameters with the aim to increase the accuracy of mutation rate estimation. The current study provides important extensions to the fluctuation data analysis using Maximum Likelihood method, with the ability to account for cell death and/or phenotypic lag, particularly with partial plating. mlemur does so in a user-friendly manner, allows to not only obtain point and interval estimates of mutation rates, but also compare them between different strains, and for many strains at the same time. mlemur also implements tools to calculate statistical power of the likelihood ratio test as well as determine the sample size to achieve prescribed power; these developments have been described in Supplementary File S1, and Table S4 contains an analysis of the required sample sizes for a wide variety of the values of *m*, which might be of help when designing a fluctuation assay.

Nevertheless, the difficulties encountered when modelling bacterial growth in the presence of a sub-inhibitory concentration of an antibiotic underline the fact that classical Luria–Delbrück distribution, even with extensions, might not be a suitable model for every fluctuation assay. Novel approaches were proposed, and might be available for the user in the future [40,42,44,70–73]. The mechanism of phenotypic lag and its impact on growth and death rate is also poorly studied, yet it has a profound impact on the distribution of mutant cells. While modern methods based on whole genome sequencing are not plagued by the same problems, the low cost and complexity as well as high quickness of generating data remain to be important advantages of the fluctuation assays, warranting further development of more sophisticated statistical models of mutation and cell proliferation.

