## Supplementary File S1 for "Efficient, robust, and versatile fluctuation data analysis using MLE MUtation Rate calculator (mlemur)"

**Supplementary File S1. Formulas for Maximum Likelihood Estimation of the number of mutations in the culture**

**for**

### Efficient, robust, and versatile fluctuation data analysis using MLE MUtation Rate calculator (mlemur)

**Krystian Łazowski[1]**

**[1] Laboratory of DNA Replication and Genome Stability, Institute of Biochemistry and Biophysics of the Polish Academy of Sciences, Pawińskiego 5a, Warsaw 02-106, Poland**

****

The probability generating function (PGF) of the Lea–Coulson distribution is:

$$G(z;m) = \exp\left[m\left(\frac{1}{z}-1\right)\log(1-z)\right]. \tag{1}$$

This PGF can be also expressed in a discretised form (Equation 18 in [1], see also [2,3]):

$$\sum\nolimits_{n=0}^{\infty} p_n z^n = \exp\Big[-m+\lambda_0+\sum\nolimits_{n=1}^{\infty}\lambda_n z^n\Big]. \tag{2}$$

We suppressed here the dependence of $m$ and $\lambda_n$ on time and tacitly assumed they are evaluated at the end of the culture growth. It is worth noting that $\lambda_n$ is also the formula for the expected number of mutations given the size of the mutant clone is $n$. Stewart et al. has established a simple protocol for calculating $\lambda_n$ and also derived the formulas for certain cases [1,4]. For the sake of future considerations, here we will re-cast the above as

$$\sum\nolimits_{n=0}^{\infty} p_n z^n = \exp\left\{m\sum_{n=0}^{\infty} h_n z^n\right\}. \tag{3}$$

As a side note, because the probability in (2) must sum to 1, $\{h_n\}$ must obviously sum to 0.

#### 1. Relaxing the perfect plating requirement in Mandelbrot–Koch distribution

Accounting for the differential growth rate of mutant and wild-type populations requires only the knowledge about their relative fitness. Following [3], the approximate PGF of M–K distribution that allows to account for differential growth is of the form

$$G(z;m,\rho) = \exp\left\{-m+\frac{m}{\rho}\sum_{k=1}^{\infty}\boldsymbol{B}\left(1;k,1+\frac{1}{\rho}\right)z^k\right\}, \tag{4}$$

where $\boldsymbol{B}$ is the (complete) beta function, and $\rho$ is the relative growth rate of a mutant cell *vs.* that of a wild-type cell. Using a more convenient parametrization $r = 1/\rho$, the values of $\boldsymbol{B}$ can calculated as given in [3]:

$$\left.\begin{aligned} B_1 &= \frac{r}{r+1} \\ B_k &= \frac{k-1}{k+r}B_{k-1} \text{ for } k \geq 2 \end{aligned}\right\}. \tag{5}$$

By rearranging (5) in the context of (3) we obtain the formula for $\{h_n\}$ when $\varepsilon = 1$:

$$\left.\begin{aligned} h_0(r) &= -1 \\ h_1(r) &= \frac{r}{r+1} \\ h_n(r) &= \frac{n-1}{n+r}h_{n-1}(r) \text{ for } n \geq 2 \end{aligned}\right\}. \tag{6}$$

When $\varepsilon \neq 1$, but $r = 1$, we can use a slightly modified version of the auxiliary sequence derived by Zheng in [5]:

$$\left.\begin{aligned} h_0(\varepsilon) &= \zeta\log\varepsilon \\ h_1(\varepsilon) &= \zeta\left(-1-\frac{\log\varepsilon}{(1-\varepsilon)}\right) \\ h_n(\varepsilon) &= \zeta\left(\frac{1}{n(n-1)}-h_{n-1}(\varepsilon)\right) \text{ for } n \geq 2 \end{aligned}\right\}, \tag{7}$$

where

$$\zeta = \frac{\varepsilon}{1-\varepsilon}. \tag{8}$$

It was discovered by Jones [6] that the recursive formula in (7) is unstable when $\varepsilon \geq 0.5$, therefore in these cases we follow Zheng [5] and calculate $\{h_n\}$ backwards by invoking the relation

$$h_n(\varepsilon) = \frac{1}{n(n+1)} - \frac{1}{\zeta} h_{n+1}(\varepsilon) \text{ for } n \geq 2. \tag{9}$$

To use this expression, however, we need to first calculate the last element of $\{h_n\}$. The exact formula for the case $\varepsilon < 1$ was derived by Stewart and Jones [4,7]:

$$\left.\begin{aligned} h_0(r,\varepsilon) &= -1 + \frac{r(1-\varepsilon)}{r+1} \boldsymbol{F}(1,1;r+2,1-\varepsilon) \\ h_n(r,\varepsilon) &= r\varepsilon^r \boldsymbol{B}(n,r+1)\, \boldsymbol{F}(r,r+1;r+1+n;1-\varepsilon) \text{ for } n \geq 1 \end{aligned}\right\}. \tag{10}$$

Here, $\boldsymbol{F}$ is the Gauss ${}_2F_1$ hypergeometric function. The strategy presented above requires computation of the value of the hypergeometric function for each element of the series, which might be time-consuming for large $n$. The computations can be facilitated using a simple relation for contiguous hypergeometric functions, given by Eq. 15.2.12 in [8]:

$$c(c-1)(z-1)\,\boldsymbol{F}(a,b;c-1;z) + c[c-1-(2c-a-b-1)z]\,\boldsymbol{F}(a,b;c;z) + (c-a)(c-b)z\,\boldsymbol{F}(a,b;c+1;z) = 0. \tag{11}$$

First, we rewrite (10) as

$$h_n(r,\varepsilon) = r\varepsilon^r B_n F_n \text{ for } n \geq 1. \tag{12}$$

$\{B_n\}$ can be calculated as given in (5). Computing $\{F_n\}$ using (11) requires calculation of two values of the hypergeometric function. To avoid the numerical instability caused by roundoff error propagation, we will follow the fashion of computing $\{F_n\}$ backwards when $\varepsilon \geq 0.5$. First, we calculate the last two elements of the $\{F_n\}$ series using (10), and then we invoke the relation

$$F_{n-1} = \frac{n(n+1)(1-\varepsilon)F_{n+1}}{\varepsilon(r+n)(r+n+1)} + \frac{[(r+n-2n(1-\varepsilon)]F_n}{\varepsilon(r+n)}. \tag{13}$$

Conversely, for $\varepsilon < 0.5$, we first compute $F_1$ and $F_2$, and then use the following formula:

$$F_{n+1} = \frac{(r+n+1)}{n(n+1)(1-\varepsilon)} \{\varepsilon(r+n)F_{n-1} - [(r+n-2n(1-\varepsilon)]F_n\}. \tag{14}$$

To summarise, the new framework requires the computation of only three values of the hypergeometric function, and three series, the third being a simple multiplication of the corresponding elements of the former two. If necessary, $\{h_n\}$ can be set to its exact value using (10).

The method of calculation of the PMF follows from the considerations of Zheng in [3,9]:

$$\left.\begin{aligned} p_0(m) &= \exp(-mh_0) \\ p_n(m) &= \frac{m}{n}\sum_{j=1}^{n} jh_j\, p_{n-j} \text{ for } n \geq 1 \end{aligned}\right\}, \tag{15}$$

$$p_n^{(1)}(m) = \{h_n\} * \{p_n(m,r,\varepsilon)\}, \tag{16}$$

$$p_n^{(2)}(m) = \{h_n\} * \left\{p_n^{(1)}(m,r,\varepsilon)\right\}. \tag{17}$$

#### 2. Accounting for phenotypic lag under the Lea–Coulson model

Phenotypic lag, or phenotypic delay, is the time required to express the selectable phenotype by cells that already acquired mutation in the marker gene. Among others, phenotypic delay was of interest to Armitage, Koch, and Angerer, and Stewart et al. [1,10–12]. The derivation of the Angerer formulation of phenotypic lag correction will be presented below.

Assuming exponential growth of bacterial cells, the number of cells $N_\tau$ at a particular time $\tau$ of culture growth is

$$N_\tau = N_0 \exp(\beta\tau), \tag{18}$$

where $N_0$ is the size of the inoculum and $\beta$ is the growth rate. The doubling time $\tau_2$ of the population is the average time required to increase the size of the population from $N_\tau$ to $2N_\tau$:

$$\tau_2 = {}^{\log(2)}\!/_{\beta}. \tag{19}$$

As argued by Stewart in [4], the probability that a mutation will occur between $t$ and $t + dt$ is $\mu N_0 \exp(\beta_1 t)\beta_1 dt$ (where $\beta_1$ is the growth rate of non-mutants), and the probability that this mutation will span to $n$ mutant cells growing until the end of the culture growth at time $t = T$ is $\exp[-\beta_2(T-t)]\{1-\exp[-\beta_2(T-t)]\}^{n-1}$ (where $\beta_2$ is the growth rate of mutants; here we assumed a pure stochastic birth of the mutant cells, see Equation 8.15 in [13]). If we assume that $\beta_1 = \beta_2 = \beta$, then the expected number of mutations that produce $n$ mutant clones between $t = 0$ and $t = T$ (which is also the number of mutations times probability that a single mutation would span to produce mutant clone of size $n$) is given by (see also Equations 46–47 in [2])

$$E[Y_n] = \int_0^T \exp[-\beta(T-t)]\{1-\exp[-\beta(T-t)]\}^{n-1}\,\mu N_0 \exp(\beta t)\beta dt. \tag{20}$$

The above equation is also a formula for $\lambda_n$ in (2). Let us now assume that the phenotypic lag is constant and equal to, say, $l$ generations. Any new mutations spanning after $t = T - l\tau_2$ will fail to produce mutant cells, which is effectively the same as if no mutations at all occurred after $T - l\tau_2$.

$\mu$ is now a function of time, but it is given by a very simple relationship:

$$\mu(t) = \begin{cases} \mu \text{ for } t \le T - l\tau_2 \\ 0 \text{ for } t > T - l\tau_2 \end{cases}.$$

We can split the time interval $[0, T]$ in (20) into $[0, T - l\tau_2]$ and $(T - l\tau_2, T]$ and zero the second integral, arriving at

$$E[Y_n] = \int_0^{T-l\tau_2} \exp[-\beta(T-t)]\{1-\exp[-\beta(T-t)]\}^{n-1}\,\mu N_0 \exp(\beta t)\beta dt. \tag{21}$$

By introducing a new variable of integration $x = \exp[-\beta(T-t)]$, $t = T + {}^{\log x}\!/_{\beta}$, $dt = {}^{dx}\!/_{\beta x}$, the upper bound of the above integral is

$$\exp\left[-\beta\left(T - T + l\frac{\log(2)}{\beta}\right)\right] = \exp[-l\log(2)] = 2^{-l} = \chi.$$

On the other hand, because the cell population size at time $T$ is $N_0\exp(\beta T)$ and at $T - l\tau_2$ it is $N_0\exp[\beta(T - l\tau_2)]$, $\chi$ is also the ratio of the two:

$$\chi = \frac{N_l}{N_t} = \frac{N_0 \exp\left[\beta\left(T - l\frac{\log(2)}{\beta}\right)\right]}{N_0\exp(\beta T)} = \exp\left[\beta T - l\beta\frac{\log(2)}{\beta} - \beta T\right] = 2^{-l}. \tag{22}$$

(We denote as $N_l$ the population size at the time when new mutations effectively stopped spanning, i.e., $T - l\tau_2$). Therefore, we can specify the phenotypic lag by the number of generations it spans, regardless of how long particular generation lasts. Coming back to (21), change of variables leads to modified Stewart's Equation 6 in [4]:

$$E[Y_n] = \mu N_t \int_0^{\chi} x^1 (1-x)^{n-1} dx = \mu N_t \boldsymbol{I}(\chi; 2, n). \tag{23}$$

This allows us to express the auxiliary sequence $\{h_n\}$ in terms of the incomplete beta function $\boldsymbol{I}(z; a, b)$:

$$h_n(\chi) = \boldsymbol{I}(\chi; 2, n) \text{ for } n \ge 1. \tag{24}$$

To find the term $h_0(\chi)$ we observe that the probability of no mutations in the culture is $e^{-m(t)}$. When there is no phenotypic lag, $-m(T) = -m$, so $h_0 = -1$ as in (6). With the phenotypic lag, if light of the constancy of mutation rate $\mu = \frac{m(T)}{N_t} = \frac{m(T - l\tau_2)}{N_l}$, $-m(T - l\tau_2) = -m\frac{N_l}{N_t} = -m\chi$, hence $h_0(\chi) = -\chi$ (see Equation 8 in [2]).

Unfortunately, for our purpose of deriving formula that accounts simultaneously for phenotypic lag and partial plating, the usage of the incomplete beta function is not practical. Instead, we will use the equation derived by Angerer in [11]:

$$\left.\begin{aligned} h_0(\Lambda) &= -\frac{1}{\Lambda} \\ h_n(\Lambda) &= \frac{1-\left(1-\frac{1}{\Lambda}\right)^n\left(1+\frac{n}{\Lambda}\right)}{n(n+1)} \text{ for } n \geq 1 \end{aligned}\right\} \tag{25}$$

where $\Lambda = 1/\chi$. It can be checked that (24) and (25) give perfectly matching results. Angerer also found the probability generating function for the distribution that induces (25):

$$G(z; m, \Lambda) = \exp\left\{m\frac{1-z}{z}\log\left(\frac{1-z}{1-z+z/\Lambda}\right)\right\}. \tag{26}$$

Following the considerations in [5], when $\varepsilon < 1$ the modified PGF assumes the form (with $\zeta$ defined in (8))

$$G(z; m, \Lambda, \varepsilon) = \exp\left\{m\zeta\frac{1-z}{1+\zeta z}\log\left(\frac{\zeta(1-z)}{\zeta(1-z)+\frac{1}{\Lambda}\left(\frac{1}{1-\varepsilon}-\zeta+\zeta z\right)}\right)\right\}. \tag{27}$$

To express (27) in the form equivalent to (3) we write $\{h_n\} = \{q_n\} * \{r_n\}$ with

$$\sum_{n=0}^{\infty} q_n z^n = \zeta(1-z)\log\left(\frac{\zeta(1-z)}{\zeta(1-z)+\frac{1}{\Lambda}\left(\frac{1}{1-\varepsilon}-\zeta+\zeta z\right)}\right), \tag{28}$$

$$\sum_{n=0}^{\infty} r_n z^n = (1+\zeta z)^{-1}. \tag{29}$$

The formula for $\{r_n\}$ has been given in [5]:

$$r_n(\varepsilon) = (-\zeta)^n \text{ for } n \geq 0. \tag{30}$$

By expanding the Taylor series in (28) at $z = 0$ it can be checked that

$$\left.\begin{aligned} q_0(\Lambda,\varepsilon) &= \zeta\log\left(\frac{-\varepsilon\Lambda}{s-1}\right) \\ q_1(\Lambda,\varepsilon) &= \zeta\left[\frac{1}{s-1}-\log\left(\frac{-\varepsilon\Lambda}{s-1}\right)\right] \\ q_n(\Lambda,\varepsilon) &= \frac{\zeta}{n(n-1)}\left[1-\frac{s^{n-1}(s-n)}{(s-1)^n}\right] \text{ for } n \geq 2 \end{aligned}\right\} \tag{31}$$

with

$$s = \varepsilon(1-\Lambda). \tag{32}$$

Finally, because (using sequence convolution)

$$h_n = (-\zeta)^n q_0 - (-\zeta)^{n-1} q_1 + \cdots + q_n,$$

$$h_{n+1} = (-\zeta)^{n+1} q_0 - (-\zeta)^n q_1 + \cdots + (-\zeta) q_n + q_{n+1},$$

we have

$$\left.\begin{aligned} h_0(\Lambda,\varepsilon) &= q_0(\Lambda,\varepsilon) \\ h_1(\Lambda,\varepsilon) &= -\zeta q_0 + q_1(\Lambda,\varepsilon) \\ h_n(\Lambda,\varepsilon) &= -\zeta h_{n-1}(\Lambda,\varepsilon) + q_n(\Lambda,\varepsilon) \text{ for } n \geq 2 \end{aligned}\right\}. \tag{33}$$

When $\varepsilon > 0.5$, $\{h_n\}$ should be calculated backwards:

$$h_n(\Lambda,\varepsilon) = -\frac{1}{\zeta}\left(h_{n+1}(\Lambda,\varepsilon) - q_{n+1}(\Lambda,\varepsilon)\right). \tag{34}$$

To find the last term $h_n$ we follow from [6] and observe that for large $n$, $h_n \approx h_{n+1}$, therefore we can write

$$h_n(\Lambda,\varepsilon) \approx -\frac{1}{\zeta}\left(h_{n(\Lambda,\varepsilon)} - q_{n+1}(\Lambda,\varepsilon)\right),$$

$$h_n(\Lambda,\varepsilon) \approx \frac{q_{n+1}(\Lambda,\varepsilon)}{\zeta+1} = (1-\varepsilon)q_{n+1}(\Lambda,\varepsilon). \tag{35}$$

Based on our trials-and-errors, it is generally sufficient to start at $q_{n+20}$ and calculate $\{h_n\}$ backwards from there.

For a stochastic phenotypic lag, we can, similarly to Crump and Hoel [14], assume that lag is constant within a clone (and distributed as above), but differing from clone to clone and drawn from Poisson distribution with mean lag $\lambda$. Obtaining $\{h_n\}$ or $\{q_n\}$ as a function of $\lambda$ is simply a matter of multiplying (25) or (31) by $P(X = l)$ with $X \sim \text{Pois}(\lambda)$ (i.e., the expression for PMF of Poisson distribution) and summing over all values of $l$:

$$\left.\begin{aligned} h_0(\lambda) &= -\frac{e^{-\lambda}}{2} \\ h_n(\lambda) &= \sum_{l=0}^{\infty}\frac{e^{-\lambda}\lambda^l}{l!}\frac{1-\left(1-\frac{1}{\Lambda}\right)^n\left(1+\frac{n}{\Lambda}\right)}{n(n+1)} \text{ for } n \geq 1 \end{aligned}\right\}, \tag{36}$$

$$\left.\begin{aligned} q_0(\lambda,\varepsilon) &= \sum_{l=0}^{\infty}\frac{e^{-\lambda}\lambda^l}{l!}\zeta\log\left(\frac{-\varepsilon l}{s-1}\right) \\ q_1(\lambda,\varepsilon) &= \sum_{l=0}^{\infty}\frac{e^{-\lambda}\lambda^l}{l!}\zeta\left[\frac{1}{s-1} - \log\left(\frac{-\varepsilon\Lambda}{s-1}\right)\right] \\ q_n(\lambda,\varepsilon) &= \sum_{l=0}^{\infty}\frac{e^{-\lambda}\lambda^l}{l!}\frac{\zeta}{n(n-1)}\left[1-\frac{s^{n-1}(s-n)}{(s-1)^n}\right] \text{ for } n \geq 2 \end{aligned}\right\}. \tag{37}$$

(36) can be expressed in terms of a finite sum involving binomial coefficients, but the algorithm is very unstable for $n > 40$ due to large values of binomial coefficient, and therefore cannot be evaluated without resorting to arbitrary precision arithmetic with hundreds of digits of precision. In practice these sequences are most efficiently evaluated by direct summation of the infinite series, which hardly ever requires summation of more than several dozen terms.

#### 3. Mutant distribution with a non-zero death rate

The equation for the exponential growth of bacterial cells is given by (18). If we assume that wild-type cells die deterministically, dying of a wild-type cell can be modelled by a process called exponential decay:

$$N_\tau = N_0\exp(-\delta\tau), \tag{38}$$

where $\delta$ is the death rate of the population. Therefore, when wild-type cells grow and die exponentially with rates, respectively, $\beta$ and $\delta$, the number of cells at time $\tau$ is given by

$$N_\tau = N_0\exp[(\beta-\delta)\tau]. \tag{39}$$

We assume $\beta > \delta$ (to ensure the population growth) and introduce $\beta^* = \beta - \delta$, the net growth rate of the population. If we express death rate as a fraction of the growth rate $\delta = d\beta$, then

$$\beta^* = \beta(1-d). \tag{40}$$

The birth and death of a cell can also be expressed in terms of a Bernoulli trial. If a living cell dies with a probability $p$ or divides into two daughter cells with probability $1-p$, then

$$\beta^* = (1-p)\beta - p\delta = (1-p)\beta - dp\beta. \tag{41}$$

By comparing (40) and (41) we find the following relations:

$$d = \frac{p}{1-p}, \tag{42}$$

$$\beta^* = \beta\frac{1-2p}{1-p}.$$

In other words, the parameter $d$ can be interpreted as the odds of dying of the cell. Because we assumed $\beta > \delta$, it follows that $d < 1$ and therefore the probability of death must be $p < 0.5$.

Equation (20) gives the expected number of mutations producing $n$ colonies assuming zero death rate. When death rate $\delta > 0$, the probability of a mutation occurring in $dt$ is $\mu N_0 \exp(\beta_1^* t)\beta_1^* dt$. On the other hand, the dynamics of the mutants can be modelled by a simple stochastic birth-and-death process [Equation 8.46 in [13], see also Equations 4–5 in [3]]:

$$\left.\begin{aligned} P_{0,\beta_2,\delta_2} &= \frac{\delta_2\left(\exp\left(\beta_2^*(T-t)\right)-1\right)}{\beta_2\exp\left(\beta_2^*(T-t)\right)-\delta_2} \\ P_{n,\beta_2,\delta_2} &= \left(1-\frac{\delta_2\left(\exp\left(\beta_2^*(T-t)\right)-1\right)}{\beta_2\exp\left(\beta_2^*(T-t)\right)-\delta_2}\right)\left(1-\frac{\beta_2\left(\exp\left(\beta_2^*(T-t)\right)-1\right)}{\beta_2\exp\left(\beta_2^*(T-t)\right)-\delta_2}\right) \\ &\qquad \left(\frac{\beta_2\left(\exp\left(\beta_2^*(T-t)\right)-1\right)}{\beta_2\exp\left(\beta_2^*(T-t)\right)-\delta_2}\right)^{n-1} \quad \text{for } n \geq 1 \end{aligned}\right\}. \tag{43}$$

We naturally define the net growth rates $\beta_1^* = \beta_1(1-d) = \beta_1[(1-2p)/(1-p)]$ and $\beta_2^* = \beta_2(1-d) = \beta_2[(1-2p)/(1-p)]$. We will only cover the case where both non-mutants and mutants have the same death rate $d$.

In the previous case we considered pure stochastic birth of the mutant cells, where it was not possible for a freshly mutated cell to produce 0 colonies assuming perfect plating; in other words, $n = 0$ mutant colonies could only result from the fact that there were 0 mutations, which we tacitly expressed as $E[Y_{n=0}] = 0$. (This should not be confused with 0 mutations occurring in a particular culture when the average number of mutations is $m$, as this probability is equal to $e^{-m}$). Now, however, there is a finite probability equal to $P_{0,\beta_2,\delta_2}$ that a mutant population has gone extinct. Hence the modified integral from (20) assumes the form

$$\left.\begin{aligned} E[Y_0] &= \int_0^T \frac{\delta_2\left(\exp\left(\beta_2^*(T-t)\right)-1\right)}{\beta_2\exp\left(\beta_2^*(T-t)\right)-\delta_2}\mu N_0 \exp(\beta_1^* t)\beta_1^* dt \\ E[Y_n] &= \int_0^T \left(1-\frac{\delta_2\left(\exp\left(\beta_2^*(T-t)\right)-1\right)}{\beta_2\exp\left(\beta_2^*(T-t)\right)-\delta_2}\right)\left(1-\frac{\beta_2\left(\exp\left(\beta_2^*(T-t)\right)-1\right)}{\beta_2\exp\left(\beta_2^*(T-t)\right)-\delta_2}\right) \\ &\qquad \left(\frac{\beta_2\left(\exp\left(\beta_2^*(T-t)\right)-1\right)}{\beta_2\exp\left(\beta_2^*(T-t)\right)-\delta_2}\right)^{n-1} \mu N_0 \exp(\beta_1^* t)\beta_1^* dt \end{aligned}\right\}. \tag{44}$$

Analogically to previous considerations, by introducing a change of variables $x = \exp\left(-\beta_2^*(T-t)\right)$ we have

$$\exp\left(\beta_2^*(T-t)\right) = \frac{1}{x};$$

$$t = T + \frac{\log(x)}{\beta_2^*};$$

$$dt = \frac{dx}{x\beta_2^*};$$

$$x(0) = \exp(-\beta_2^* T) = \left(\frac{N_0}{N_t}\right)^{\rho} \approx 0;$$

$$x(T) = 1;$$

$$\exp\left(\beta_1^*\left(T + \frac{\log(x)}{\beta_2^*}\right)\right) = \left(\frac{N_t}{N_0}\right)x^r.$$

Substituting into (44) and after simple algebra we obtain

$$\left.\begin{aligned} E[Y_0] &= \mu N_t r d \int_0^1 \frac{(1-x)x^{r-1}}{1-d\,x} dx, \\ E[Y_n] &= \mu N_t r (1-d)^2 \int_0^1 \frac{(1-x)^{n-1}x^r}{(1-d\,x)^{n+1}} dx \ \text{ for } n \geq 1 \end{aligned}\right\}. \tag{45}$$

The integrals can be re-cast in terms of the product of beta and hypergeometric functions (see Equation 15.3.1 in [8]), which leads us to the solution:

$$\left.\begin{aligned} h_0(r,d) &= -1 + rd\,\boldsymbol{B}(r;2)\,\boldsymbol{F}(1,r;r+2;d) \\ h_n(r,d) &= (1-d)^2 r\,\boldsymbol{B}(n;1+r)\,\boldsymbol{F}(n+1,r+1;r+1;d) \text{ for } n \geq 1 \end{aligned}\right\}. \tag{46}$$

To allow $\varepsilon < 1$ let us assume that each of the mutant cells has the probability $\varepsilon$ of growing on the selective medium and the probability $1-\varepsilon$ of not being observed. Hence the probability that of $n = k + m$ mutants, $k$ will be observed while $m$ will not is given by binomial distribution:

$$P_{k,l} = \binom{k+m}{k} \varepsilon^k (1-\varepsilon)^m. \tag{47}$$

By marginalising over $m$ we can derive the expected number of mutations when $k$ mutants are observed. When $k = 0$, setting $m = 0$ gives $n = 0$ which is the case given by $E[Y_0]$, and setting $m > 0$ gives $n = m$ which is given by $E[Y_m]$, therefore (we continue employing Stewart's notation from [4] and denote the expected number of mutations that between $t = 0$ and $t = T$ produced $k \leq n$ observed mutants as $E[Z_0]$)

$$E[Z_0] = E[Y_0] + \sum_{m=1}^{\infty} (1-\varepsilon)^m E[Y_m] = E[Y_0] + \mu N_t r (1-d)^2 \sum_{m=1}^{\infty} \left( \int_0^1 (1-\varepsilon)^m \frac{(1-x)^{m-1}}{(1-d\,x)^{m+1}} x^r dx \right). \tag{48}$$

The integral is given by the hypergeometric function but evaluating infinite sum of the hypergeometric functions can pose a significant computational problem. Instead, we observe that the expression under the integral is strictly non-negative because $x \in [0,1]$ and $\varepsilon, d \in (0,1)$, hence we can interchange the order of summation and integration:

$$E[Z_0] = E[Y_0] + \mu N_t r (1-d)^2 \int_0^1 x^r \left( \sum_{m=1}^{\infty} (1-\varepsilon)^m \frac{(1-x)^{m-1}}{(1-d\,x)^{m+1}} \right) dx.$$

The infinite sum $\sum_{m=1}^{\infty} (1-\varepsilon)^m \frac{(1-x)^{m-1}}{(1-d\,x)^{m+1}}$ is a geometric series $\sum_{n=1}^{\infty} a q^{n-1}$ with

$$a = \frac{1-\varepsilon}{(1-d\,x)^2},$$
$$q = \frac{(1-\varepsilon)(1-x)}{(1-d\,x)}.$$

The series converges when $|q| < 1$, which is always true for $x \in [0,1]$ and $\varepsilon, d \in (0,1)$. The proof is very simple. Setting $x = 0$ gives $q = 1 - \varepsilon < 1$. Setting $x = 1$ gives $q = 0$. When $x \in (0,1)$ assume that the above is true. Then

$$-1 < \frac{(1-\varepsilon)(1-x)}{(1-dx)} < 1$$
$$dx - 1 < 1 - \varepsilon - x + \varepsilon x < 1 - dx$$
$$x(d+1) - 2 < \varepsilon(x-1) < x(1-d)$$
$$\frac{x(d+1)-2}{x-1} > \varepsilon > \frac{x(1-d)}{x-1}.$$

Because $x - 1 < 0$ the right-hand side must be true. Observe that

$$\frac{x(d+1)-2}{x-1} = \frac{2-x(d+1)}{1-x} = \frac{1+1-x\,d-x}{1-x} = \frac{1-x\,d}{1-x} + 1$$

which is always bigger than 1 and therefore always bigger than $\varepsilon$.

From the elementary properties of the geometric series

$$\lim_{n\to\infty} a \frac{1-q^n}{1-q} = \frac{a}{1-q} = \frac{(1-\varepsilon)}{(1-d\,x)[\varepsilon(1-x) + x(1-d)]}.$$

Therefore, after partial fraction expansion we arrive at

$$E[Z_0] = E[Y_0] + \mu N_t r(1-d)\left\{\int_0^1 \frac{1}{\varepsilon}\frac{(1-d-\varepsilon)x^r}{1-x\left(\frac{d+\varepsilon-1}{\varepsilon}\right)}dx + \int_0^1 \frac{d\,x^r}{(1-d\,x)}dx\right\}. \quad (49)$$

Both integrals are given by the hypergeometric function. We need to note, however, that additional caution must be exercised when evaluating the first term, depending on the value of $d$. The hypergeometric function converges only when $\left|{}^{d+\varepsilon-1}/_{\varepsilon}\right| < 1$, which is the case when $d > 1-2\varepsilon$. When $d \leq 1-2\varepsilon$ we can use one of the linear transformation formulas (i.e., 15.3.4 or 15.3.5 from [8]) that impose a new radius of convergence $\left|{}^{d+\varepsilon-1}/_{d-1}\right| < 1$, which implies $d < 1-{}^{\varepsilon}/_{2}$. Because when $d$ is $\leq 1-2\varepsilon$ (which violates the former case) it cannot be $> 1-{}^{\varepsilon}/_{2}$ (which would violate the latter case), these two cases cover the whole allowed parameter space. To summarise:

$$E[Z_0] = E[Y_0] + \mu N_t r\frac{1-d}{r+1}\left[d\boldsymbol{F}(1,1+r;2+r;d) + \frac{(1-\varepsilon-d)}{\varepsilon}\left\{\begin{matrix} \boldsymbol{F}\left(1,1+r;2+r;\frac{\varepsilon+d-1}{\varepsilon}\right) & d > 1-2\varepsilon \\ \frac{\varepsilon}{1-d}\boldsymbol{F}\left(1,1;2+r;\frac{\varepsilon+d-1}{d-1}\right) & \text{otherwise}\end{matrix}\right\}\right]. \quad (50)$$

We will now treat the case when $k > 0$, which implies $n > 0$. Subsequent steps of derivation are similar:

$$\begin{aligned}
E[Z_k] &= \sum_{m=0}^{\infty}\binom{k+m}{k}\varepsilon^k(1-\varepsilon)^m E[Y_{k+m}] = \mu N_t r(1-d)^2\sum_{m=0}^{\infty}\left(\int_0^1\binom{k+m}{k}\varepsilon^k(1-\varepsilon)^m\frac{(1-x)^{k+m-1}x^r}{(1-d\,x)^{k+m+1}}dx\right) \\
&= \mu N_t r(1-d)^2\int_0^1\frac{(1-x)^{k-1}x^r}{(1-d\,x)^{k+1}}\varepsilon^k\left(\sum_{m=0}^{\infty}\binom{k+m}{k}(1-\varepsilon)^m\frac{(1-x)^m}{(1-d\,x\,)^m}\right)dx \\
&= \mu N_t\frac{r(1-d)^2}{\varepsilon}\int_0^1\frac{(1-x)^{k-1}x^r}{\left(1-x\frac{(\varepsilon+d-1)}{\varepsilon}\right)^{k+1}}dx \\
&= \mu N_t\frac{r(1-d)^2}{\varepsilon}\boldsymbol{B}(k;1+r)\left\{\begin{matrix}\boldsymbol{F}\left(k+1,r+1;k+r+1;\frac{\varepsilon+d-1}{\varepsilon}\right) & d > 1-2\varepsilon \\ \left(\frac{\varepsilon}{1-d}\right)^{r+1}\boldsymbol{F}\left(r+1,r;r+1;\frac{\varepsilon+d-1}{d-1}\right) & \text{otherwise}\end{matrix}\right\}.
\end{aligned} \quad (51)$$

Consequently, the formulas for $\{h_n\}$ are given by

$$\left.\begin{aligned}
h_0(r,d,\varepsilon) &= -1 + rd\boldsymbol{B}(r^*;2)\boldsymbol{F}(1,r;r+2;d) \\
&\quad + \frac{1-d}{r+1}\left[d\boldsymbol{F}(1,1+r;2+r;d) + \frac{(1-\varepsilon-d)}{\varepsilon}\left\{\begin{matrix}\boldsymbol{F}\left(1,1+r;2+r;\frac{\varepsilon+d-1}{\varepsilon}\right) & d > 1-2\varepsilon \\ \frac{\varepsilon}{1-d}\boldsymbol{F}\left(1,1;2+r;\frac{\varepsilon+d-1}{d-1}\right) & \text{otherwise}\end{matrix}\right\}\right] \\
h_n(r,d,\varepsilon) &= \frac{r(1-d)^2}{\varepsilon}\boldsymbol{B}(k;1+r)\left\{\begin{matrix}\boldsymbol{F}\left(n+1,r+1;n+r+1;\frac{\varepsilon+d-1}{\varepsilon}\right) & d > 1-2\varepsilon \\ \left(\frac{\varepsilon}{1-d}\right)^{\text{r}+1}\boldsymbol{F}\left(r+1,r;n+r+1;\frac{\varepsilon+d-1}{d-1}\right) & \text{otherwise}\end{matrix}\right\} \text{ for } n \geq 1
\end{aligned}\right\}. \quad (52)$$

#### 4. Extending the gamma mixture method to the special cases

The usage of Luria–Delbrück distribution requires that the final number of cells is identical (or, in practice, at least reasonably similar) in each test tube. The influence of the variability of $N_t$ on MLE of $m$ has been reviewed in [15], and the most relevant conclusions will be presented below.

First, we can express the variability of $N_t$ by its coefficient of variation ($C_V$), which is a standardised measure of dispersion. Further, we assume that the number of mutations in each culture, $m_i$, is proportional to the total number of cells in the same tube, $N_{t,i}$. Therefore, the coefficient of variation of $m$ is expected to be the same as the coefficient of variation of $N_t$ (Eq. 11 in [15]).

A convenient and reliable way of accounting for the variability in the final number of cells in bacterial cultures is to use the gamma mixture method described in [15]. In this method, it is assumed that the mutant counts obey a gamma–LD distribution, i.e., a $B^0$ distribution indexed by the following parameters:

$$k = C_V{}^{-2}, \tag{53}$$

$$A = \frac{m}{k} = mC_V{}^2. \tag{54}$$

The probability generating function (PGF) of the $B^0$ distribution has been given in [16]:

$$G(z; A, k) = \exp\left\{-k\log\left[1 - A\left(\frac{1}{z} - 1\right)\log(1-z)\right]\right\}. \tag{55}$$

By using the similarities between (1) and (55), in light of (3), we can recast the above as

$$G(z; m, k) = \exp\left\{-k\log\left[1 - \frac{m}{k}\sum_{n=0}^{\infty} h_n z^n\right]\right\}. \tag{56}$$

When expressed like this, the gamma–LD mixture distribution can readily utilise previously derived $\{h_n\}$, which essentially relaxes all the requirements for which the formula for $\{h_n\}$ has been found.

We now need to temporarily introduce another auxiliary sequence:

$$\sum_{n=0}^{\infty} \vartheta_n z^n = 1 - \frac{m}{k}\sum_{n=0}^{\infty} h_n z^n. \tag{57}$$

Calculating $\{\vartheta_n\}$ is trivial:

$$\left.\begin{aligned} \vartheta_0 &= 1 - \frac{m}{k} h_0 \\ \vartheta_n &= -\frac{m}{k} h_n \text{ for } n \geq 1 \end{aligned}\right\}. \tag{58}$$

Now we can directly apply Lemmas 1 and 2 from [16]. First, we observe that

$$\left.\begin{aligned} \xi_0 &= \log(\vartheta_0) = \log\left(1 - \frac{m}{k}h_0\right) \\ \xi_n &= \frac{1}{n\left(1 - \frac{m}{k}h_0\right)}\left(n\vartheta_n - \sum_{j=1}^{n-1} j\xi_j \vartheta_{n-j}\right) \text{ for } n \geq 1 \\ \xi_n &= \frac{-m/k}{n\left(1 - \frac{m}{k}h_0\right)}\left(nh_n - \sum_{j=1}^{n-1} j\xi_j h_{n-j}\right) \text{ for } n \geq 1 \end{aligned}\right\}. \tag{59}$$

Second, to calculate the probability mass function, we have

$$\left.\begin{aligned} p_0(m,k) &= \exp(-k\xi_0) = \left(1 - \frac{m}{k}h_0\right)^{-k} \\ p_n(m,k) &= -\frac{k}{n}\sum_{j=1}^{n} j\xi_j p_{n-j} \text{ for } n \geq 1 \end{aligned}\right\}. \tag{60}$$

We can now calculate the first and second derivatives of the PGF given by (56). For clarity, we shall substitute $\{h_n\}$ given by (10) into (56).

$$\begin{aligned} \frac{\partial G(z; m, k)}{\partial m} &= \exp\left[-k\log\left(1 - \frac{m}{k}\{h_n\}\right)\right] \cdot (-k) \cdot \left[\left(1 - \frac{m}{k}\{h_n\}\right)^{-1}\right] \cdot \left(-\frac{\{h_n\}}{k}\right) \\ &= \exp[-(k+1)\log(1 - A\zeta\{h_n\})] \cdot \{h_n\} = \{h_n\}G(z; m, k+1), \end{aligned} \tag{61}$$

$$\frac{\partial^2}{\partial m^2}G(z; m, k) = \frac{\partial}{\partial m}\left(\frac{\partial G(z; m, k)}{\partial m}\right) = \{h_n\}\frac{\partial}{\partial m}G(z; m, k+1) = \frac{(k+1)}{k}\{h_n\}\{h_n\}G(z; m, k+2). \tag{62}$$

From this we can conclude the forms of the first and second derivatives of $p$ (with $*$ denoting sequence convolution):

$$p_n^{(1)}(m,k) = \{h_n\} * \{p_n(m,k+1)\}, \tag{63}$$

$$p_n^{(2)}(m,k) = \frac{(k+1)}{k}\{h_n\}^{*2} * \{p_n(m,k+2)\}. \tag{64}$$

MLE of $m$ can be calculated using an iterative Newton algorithm that involves first and second derivatives of the log-likelihood function. Confidence intervals and statistical significance of the difference between two mutation rates can be calculated using likelihood ratio tests. All necessary formulas were given in [16,17], but for the reader's convenience will also be given in Section 6.

#### 5. A universal sequence

In light of the equations developed in this work it is possible to derive a general integral representation of $\{h_n\}$ that combines corrections for: (a) size of the inoculum; (b) differential growth rate of mutants and non–mutants; (c) cell death; (d) phenotypic lag; (e) partial plating. The formulas will be presented here for educational purposes. Apart from the previously used symbols, we here define $\varphi = {}^{N_0}\!/_{N_t}$. These formulas are universal in the sense that they can replace any other expression for $\{h_n\}$; however, it does not mean that all corrections may be used together at the same time. For reasons explained in the paper we discourage setting $r \neq 1$ together with $\chi < 1$.

When $d = 0$ and $\varepsilon = 1$

$$\left.\begin{aligned} &h_0(\varphi, r, d=0, \chi, \varepsilon=1) = -\chi^r, \\ &h_n(\varphi, r, d=0, \chi, \varepsilon=1) = (1-\varphi\chi^{-r})^{-1}\int_{\varphi^{1/r}}^{\chi} x^r(1-x)^{n-1}dx \end{aligned}\right\}. \tag{65}$$

Otherwise

$$\left.\begin{aligned} &h_0(\varphi, r, d, \chi, \varepsilon) = -\chi^r + (1-\varphi\chi^{-r})^{-1}\int_{\varphi^{1/r}}^{\chi}\left\{\frac{rd(1-x)x^{r-1}}{1-d\,x} + (1-d)r\left[\frac{(1-d-\varepsilon)x^r}{\varepsilon - x(d+\varepsilon-1)} + \frac{dx^r}{1-d\,x}\right]\right\}dx \\ &h_n(\varphi, r, d, \chi, \varepsilon) = (1-\varphi\chi^{-r})^{-1}r(1-d)^2\int_{\varphi^{1/r}}^{\chi}\frac{(1-x)^{n-1}x^r}{(1-d\,x)^{n+1}}dx \end{aligned}\right\}. \tag{66}$$

These formulas can be evaluated using numerical integration. In our experience tanh-sinh quadrature worked well in all tested cases.

#### 6. Point and interval estimates of *m* using Newton-Raphson algorithm

To find MLE of $m$, we iteratively solve the following equation to a satisfactory degree of accuracy:

$$m_{k+1} = m_k + \frac{U(m_k)}{J(m_k)}. \tag{67}$$

$U(m_k)$ is called the score function and is a first derivative of the log-likelihood function:

$$l(m) = \sum\nolimits_{i=1}^{n}\log\left[p(X_i;m)\right], \tag{68}$$

$$U(m) = \frac{\partial l(m)}{\partial m} = \frac{\partial p(m)}{\partial m}\frac{1}{p(m)} = \sum\nolimits_{i=1}^{n}\frac{p^{(1)}(X_i;m)}{p(X_i;m)}. \tag{69}$$

The observed Fisher information $J(m_k)$ is the negative second derivative of the log-likelihood function:

$$\begin{aligned} J(m) &= -\frac{\partial^2 l(m)}{\partial m^2} = -\frac{\partial}{\partial m}\left(\frac{\partial p(m)}{\partial m}\frac{1}{p(m)}\right) = \left(\frac{\partial p(m)}{\partial m}\right)^2\frac{1}{p(m)^2} - \frac{\partial^2 p(m)}{\partial m^2}\frac{1}{p(m)} \\ &= \sum\nolimits_{i=1}^{n}\left[\left(\frac{p^{(1)}(X_i;m)}{p(X_i;m)}\right)^2 - \frac{p^{(2)}(X_i;m)}{p(X_i;m)}\right]. \end{aligned} \tag{70}$$

Here, $X_i$ is a single mutant count, hence we sum those values of the PMF (or its derivatives) that correspond to the obtained mutant counts. The formulas for the probabilities, as well as their first and second derivatives, are given by (15), (16), (17), (60), (63), and (64). Equations (67) – (70) are universal in the sense that they can be used regardless of the underlying assumptions.

Confidence intervals for $m$ can be constructed using inverted likelihood ratio test. According to Wilks' theorem, twice negative logarithm of the likelihood ratio statistic is asymptotically chi-squared-distributed with 1 degree of freedom:

$$-2\log\lambda = -2[l(\widehat{m}_\alpha) - l(\widehat{m})] \approx \chi_1^2(\alpha), \tag{71}$$

from which it follows that that at the boundary of the $100(1-\alpha)\%$ confidence interval $m_\alpha$ assumes one of two possible values such that

$$l(\widehat{m}_\alpha) = l(\widehat{m}) - \frac{1}{2}\chi_1^2(\alpha). \tag{72}$$

To find $m_\alpha$, we iteratively solve the following equation:

$$m_{\alpha,k+1} = m_{\alpha,k} - \frac{l(m_\alpha)}{U(m_{\alpha,k})}. \tag{73}$$

Because (73) has two solutions, to ensure proper convergence it is imperative to choose correct initial value $m_{\alpha,0}$. One way to do this is to set $m_{\alpha,0}$ close to one of the theoretical roots assuming that log-likelihood is quadratic at $\widehat{m}$. Therefore

$$m_{\alpha,0} = \widehat{m} \pm \frac{1}{2}\sqrt{\frac{\chi_1^2(\alpha)}{J(\widehat{m})}}. \tag{74}$$

The above method may fail when the distribution is extremely heavily tailed, for example when mutant cells grow much faster that non-mutants or when coefficient of variation of the number of cells in culture is big. However, even then a satisfactory starting value can usually be found by trial-and-error. Alternatively, one can use a hybrid bisection-Newton algorithm which is usually more stable.

#### 7. Power analysis and sample size determination for likelihood ratio test

To find the formulas for power and determining sample size for likelihood ratio test we follow the approach described in [18]. The problem under consideration is analogical to comparing two mutation rates as previously tackled by Zheng [17]. In essence, we have two datasets with mutation rates $\mu_1 \neq \mu_2$ along with culture sizes $N_{t1}, N_{t2}$ and sample sizes $n_1, n_2$. We want to calculate the statistical power of the likelihood ratio test, which is the probability that the differences in mutation rates will be statistically significant, or in other words the probability that the likelihood ratio statistic $\lambda$ will be bigger than $(1-\alpha)$ quantile of the chi-squared distribution with one degree of freedom (which is the distribution of the LR statistic under the null hypothesis):

$$power = P\big(\lambda > \chi_1^2(\alpha)\big). \tag{75}$$

For the level of significance $\alpha = 0.05$, $\chi_1^2(\alpha) \approx 3.841$. Alternatively, we want to calculate the required sample size $n = n_1 = n_2$ to achieve a prescribed level of statistical power. In any case, under certain regularity conditions, the distribution of the likelihood ratio statistic under the alternative hypothesis is a non-central chi-squared distribution with some non-centrality parameter $D$ and one degree of freedom, that is, $\lambda \sim \chi_{1,D}^2$ [19]. The non-centrality parameter is equal to twice negative difference between conditional expectations of the log-likelihood function under the null and alternative hypotheses for a single observation, times sample size, assuming that the alternative hypothesis is true [18]. When dealing with two samples, $D$ is simply a linear combination of $D_1$ and $D_2$. The exact formula, modified to our case, is therefore

$$D = 2\left\{n_1\big(\mathbb{E}[l_{m_1}(m_1)] - \mathbb{E}[l_{m_1}(\widetilde{m}_C)]\big) + n_2\left(\mathbb{E}[l_{m_2}(m_2)] - \mathbb{E}\left[l_{m_2}\left(\frac{N_{t1}}{N_{t2}}\widetilde{m}_C\right)\right]\right)\right\}. \tag{76}$$

$\widetilde{m}_C$ is MLE of $m$ under the null hypothesis, that is, when we assume that $\mu_1 = \mu_2 = \mu$. In other words, one dataset is LD–distributed with parameter $\widetilde{m}_C$ while the other with parameter $(N_{t1}/N_{t2})\widetilde{m}_C$. This value can be estimated as given in [17] by simulating and pooling large-sample fluctuation data for $\mu_1, \mu_2$ and $N_{t1}, N_{t2}$, but because probability is by (frequentist) definition a limiting frequency of an outcome given infinitely many trials, $\tilde{m}_C$ can be also found directly using the known values of probability mass function for $m_1$ and $m_2$, $p(m_1)$ and $p(m_2)$:

$$\tilde{m}_C = \underset{m}{\operatorname{argmax}} \left\{ \sum_{i=0}^{\infty} \left[ l(m, p_i(m_1)) + l\left(\frac{N_{t_1}}{N_{t_2}} m, p_i(m_2)\right) \right] \right\}. \tag{77}$$

$\tilde{m}_C$ can be computed using an algorithm analogical to the one described in section 6, but using conditional expectations of score and Fisher information:

$$U_C(m) = \frac{\partial l_C(m)}{\partial m} = \sum_{i=0}^{\infty} \frac{p_i^{(1)}(m)}{p_i(m)} p_i(m_1) + \frac{N_{t_1}}{N_{t_2}} \sum_{i=0}^{\infty} \frac{p_i^{(1)}(m)}{p_i(m)} p_i(m_2), \tag{78}$$

$$J_C(m) = -\frac{\partial^2 l_C(m)}{\partial m^2} = \sum_{i=0}^{\infty} \left[ \left(\frac{p_i^{(1)}(m)}{p_i(m)}\right)^2 - \frac{p_i^{(2)}(m)}{p_i(m)} \right] p_i(m_1) + \left(\frac{N_{t_1}}{N_{t_2}}\right)^2 \sum_{i=0}^{\infty} \left[ \left(\frac{p_i^{(1)}(m)}{p_i(m)}\right)^2 - \frac{p_i^{(2)}(m)}{p_i(m)} \right] p_i(m_2). \tag{79}$$

Then

$$\mathbb{E}[l_{m_1}(m_1)] = \sum_{i=0}^{\infty} l_i(m_1) p_i(m_1), \tag{80}$$

$$\mathbb{E}[l_{m_2}(m_2)] = \sum_{i=0}^{\infty} l_i(m_2) p_i(m_2), \tag{81}$$

$$\mathbb{E}[l_{m_1}(\tilde{m}_C)] = \sum_{i=0}^{\infty} l_i(\tilde{m}_C) p_i(m_1), \tag{82}$$

$$\mathbb{E}\left[l_{m_2}\left(\frac{N_{t_1}}{N_{t_2}} \tilde{m}_C\right)\right] = \sum_{i=0}^{\infty} l_i\left(\frac{N_{t_1}}{N_{t_2}} \tilde{m}_C\right) p_i(m_2). \tag{83}$$

The infinite series can usually be truncated at several thousand terms depending on the values of $m$, such that CDF is sufficiently close to unity. Once the value of the non-centrality parameter has been obtained, statistical power is found using the value of cumulative distribution function of $\chi^2_{1,D}(x)$ where $x = \chi^2_1(\alpha)$, that is, $1 - F(\chi^2_1(\alpha); 1d.f., D)$. Conversely, the required sample size to obtain a prescribed statistical power is calculated by finding $n$ that gives $D$ such that $F(\chi^2_1(\alpha); 1d.f., D) = 1 - power$. For a typical recommended power of 80%, the corresponding value of $D$ is ~7.849.
