## Supplementary File S2 for "Efficient, robust, and versatile fluctuation data analysis using MLE MUtation Rate calculator (mlemur)"

**Supplementary File S2. Supplementary Tables and Figures**

**for**

### Efficient, robust, and versatile fluctuation data analysis using MLE MUtation Rate calculator (mlemur)

**Krystian Łazowski[1]**

**[1] Laboratory of DNA Replication and Genome Stability, Institute of Biochemistry and Biophysics of the Polish Academy of Sciences, Pawińskiego 5a, Warsaw 02-106, Poland**

****

**Table S1. The usefulness of stochastic Angerer formulation of the Luria–Delbrück distribution against protein dilution-simulated data with the threshold value 0.**

The threshold value of wild-type protein units in the cell above which the phenotype is not expressed has been set to 0. $N$ — number of experiments simulated $n$ — number of cultures in each experiment; $m$ — average number of mutations; $u$ — initial number of protein units; $\lambda$ — average phenotypic lag. Joint estimation — MLEs of $m$ and $\lambda$ were estimated simultaneously under the stochastic Angerer model. LC formulation — estimation of m under the Lea–Coulson formulation. Nominal CI coverage is 95%.

| **Simulation parameters** | | | | **Joint estimation** | | | **LC formulation** | |
|---|---|---|---|---|---|---|---|---|
| $N$ | $n$ | $m$ | $u$ | median $m$ | CI coverage | median $\lambda$ | median $m$ | CI coverage |
| 10000 | 100 | 1 | 2 | 0.930 | 94.4% | 1.41 | 0.486 | 0.2% |
| 10000 | 100 | 1 | 4 | 0.900 | 94.2% | 1.95 | 0.359 | 0.0% |
| 10000 | 100 | 1 | 8 | 0.861 | 94.4% | 2.52 | 0.259 | 0.0% |
| 10000 | 100 | 1 | 32 | 0.709 | 92.3% | 3.35 | 0.135 | 0.0% |
| 10000 | 100 | 10 | 2 | 9.38 | 93.0% | 1.45 | 6.12 | 0.0% |
| 10000 | 100 | 10 | 4 | 9.17 | 92.2% | 2.03 | 4.78 | 0.0% |
| 10000 | 100 | 10 | 8 | 8.87 | 91.0% | 2.59 | 3.58 | 0.0% |
| 10000 | 100 | 10 | 32 | 7.76 | 83.5% | 3.54 | 1.85 | 0.0% |
| 10000 | 30 | 1 | 2 | 0.908 | 96.3% | 1.36 | 0.485 | 21.6% |
| 10000 | 30 | 1 | 4 | 0.875 | 95.0% | 1.89 | 0.355 | 4.7% |
| 10000 | 30 | 1 | 8 | 0.833 | 94.6% | 2.42 | 0.253 | 0.8% |
| 10000 | 30 | 1 | 32 | 0.675 | 95.1% | 3.19 | 0.143 | 0.0% |
| 10000 | 30 | 10 | 2 | 9.07 | 93.9% | 1.31 | 6.15 | 4.5% |
| 10000 | 30 | 10 | 4 | 8.97 | 93.3% | 1.94 | 4.82 | 0.3% |
| 10000 | 30 | 10 | 8 | 8.61 | 92.6% | 2.49 | 3.60 | 0.0% |
| 10000 | 30 | 10 | 32 | 7.56 | 90.1% | 3.48 | 1.85 | 0.0% |

**Table S2. Coefficients of variation of the deaths-to-births ratio in data used in Table 6.**

$N_E$ — number of experiments simulated; $n_C$ — number of cultures in each experiment; $N_0$ — size of inoculum; $t$ — time of culture growth; $\mu$ — mutation rate per cell; $d$ — relative death rate; $\rho$ — relative mutant fitness; $N_t + n$ — average final number of cells in the culture (non-mutant and mutant).

| Simulation parameters | | | | | | | | Deaths/Births *CV* |
|---|---|---|---|---|---|---|---|---|
| Line | $N_E$ | $n_C$ | $N_0$ | $t$ | $\mu$ | $d$ | $\rho$ | |
| 1 | 10000 | 100 | 100 | 9.2 | $1{\times}10^{-4}$ | 0.5 | 1 | 1.3% |
| 2 | 10000 | 100 | 100 | 12.4 | $1{\times}10^{-4}$ | 0.5 | 1 | 0.56% |
| 3 | 10000 | 100 | 100 | 13.8 | $1{\times}10^{-4}$ | 0.5 | 1 | 0.39% |
| 4 | 10000 | 100 | 200 | 12.4 | $1{\times}10^{-4}$ | 0.5 | 1 | 0.39% |
| 5 | 10000 | 100 | 100 | 15.4 | $1{\times}10^{-4}$ | 0.7 | 1 | 0.88% |
| 6 | 10000 | 100 | 100 | 20.7 | $1{\times}10^{-4}$ | 0.7 | 1 | 0.39% |
| 7 | 10000 | 100 | 100 | 20.7 | $1{\times}10^{-4}$ | 0.7 | 0.8 | 0.39% |
| 8 | 10000 | 100 | 100 | 23 | $1{\times}10^{-4}$ | 0.7 | 1 | 0.28% |
| 9 | 10000 | 100 | 100 | 23 | $1{\times}10^{-5}$ | 0.7 | 1 | 0.28% |
| 10 | 10000 | 100 | 250 | 17.7 | $1{\times}10^{-4}$ | 0.7 | 1 | 0.39% |
| 11 | 10000 | 100 | 250 | 20 | $1{\times}10^{-4}$ | 0.7 | 1 | 0.27% |
| 12 | 10000 | 100 | 500 | 17.7 | $1{\times}10^{-4}$ | 0.7 | 1 | 0.27% |
| 13 | 10000 | 100 | 250 | 53 | $1{\times}10^{-4}$ | 0.9 | 1 | 0.22% |
| 14 | 10000 | 100 | 500 | 46.1 | $1{\times}10^{-4}$ | 0.9 | 1 | 0.21% |
| 15 | 10000 | 100 | 500 | 53 | $1{\times}10^{-4}$ | 0.9 | 1 | 0.15% |
| 16 | 10000 | 100 | 500 | 53 | $1{\times}10^{-5}$ | 0.9 | 1 | 0.15% |
| 17 | 10000 | 100 | 1000 | 46.1 | $1{\times}10^{-4}$ | 0.9 | 1 | 0.15% |
| 18 | 10000 | 100 | 1000 | 55.2 | $1{\times}10^{-4}$ | 0.9 | 1 | 0.093% |
| 19 | 10000 | 100 | 2000 | 39.1 | $1{\times}10^{-4}$ | 0.9 | 1 | 0.15% |

**Table S3. Point and interval estimates of $\mu$ under a fully stochastic model with cell death using data from Table 6, but for $n_C$ = 30.**

$N_E$ — number of experiments simulated; $n_C$ — number of cultures in each experiment; $N_0$ — size of inoculum; $t$ — time of culture growth; $\mu$ — mutation rate per cell; $d$ — relative death rate; $\rho$ — relative mutant fitness; $N_t + n$ — average final number of cells in the culture (non-mutant and mutant). The same simulated data as in Table 6 were used here, but only first 30 cultures from each dataset were used. The data were modelled using a simple birth-death process for mutants with deterministic growth of wild-types, with consideration for the inoculum, using either Luria–Delbrück distribution or $B^0$ distribution where $CV$ is taken into account. For each experiment, $\mu$ was calculated by dividing the estimate of $m$ by $(N_t + n - N_0)/(1 - d)$. Mean and $CV$ for $N_t$ was calculated for each experiment, and median values are presented in the table. Nominal CI coverage is 95%.

| | Simulation parameters | | | | | | | Culture size | | LD distribution | | $B^0$ distribution | | Golden Benchmark | |
|---|---|---|---|---|---|---|---|---|---|---|---|---|---|---|---|
| Line | $N_E$ | $n_C$ | $N_0$ | $t$ | $\mu$ | $d$ | $\rho$ | median $N_t + n$ | median $CV$ | median $\mu$ | CI coverage | median $\mu$ | CI coverage | median $\mu$ | CI coverage |
| 1 | 10000 | 30 | 100 | 9.2 | $1{\times}10^{-4}$ | 0.5 | 1 | $9.95{\times}10^{3}$ | 17.0% | $9.87{\times}10^{-5}$ | 95.2% | $9.99{\times}10^{-5}$ | 95.4% | $9.98{\times}10^{-5}$ | 95.2% |
| 2 | 10000 | 30 | 100 | 12.4 | $1{\times}10^{-4}$ | 0.5 | 1 | $4.93{\times}10^{4}$ | 17.1% | $9.79{\times}10^{-5}$ | 94.3% | $1.01{\times}10^{-4}$ | 95.6% | $1.01{\times}10^{-4}$ | 94.9% |
| 3 | 10000 | 30 | 100 | 13.8 | $1{\times}10^{-4}$ | 0.5 | 1 | $9.93{\times}10^{4}$ | 17.1% | $9.73{\times}10^{-5}$ | 93.2% | $1.00{\times}10^{-4}$ | 95.9% | $1.00{\times}10^{-4}$ | 94.9% |
| 4 | 10000 | 30 | 200 | 12.4 | $1{\times}10^{-4}$ | 0.5 | 1 | $9.87{\times}10^{4}$ | 12.1% | $9.89{\times}10^{-5}$ | 94.3% | $1.00{\times}10^{-4}$ | 95.3% | $1.00{\times}10^{-4}$ | 95.0% |
| 5 | 10000 | 30 | 100 | 15.4 | $1{\times}10^{-4}$ | 0.7 | 1 | $1.02{\times}10^{4}$ | 23.4% | $9.81{\times}10^{-5}$ | 95.1% | $1.01{\times}10^{-4}$ | 95.5% | $1.01{\times}10^{-4}$ | 95.0% |
| 6 | 10000 | 30 | 100 | 20.7 | $1{\times}10^{-4}$ | 0.7 | 1 | $4.98{\times}10^{4}$ | 23.5% | $9.56{\times}10^{-5}$ | 92.8% | $1.01{\times}10^{-4}$ | 96.3% | $1.01{\times}10^{-4}$ | 95.1% |
| 7 | 10000 | 30 | 100 | 20.7 | $1{\times}10^{-4}$ | 0.7 | 0.8 | $4.98{\times}10^{4}$ | 23.6% | $9.55{\times}10^{-5}$ | 92.1% | $1.00{\times}10^{-4}$ | 96.2% | $1.00{\times}10^{-4}$ | 94.8% |
| 8 | 10000 | 30 | 100 | 23 | $1{\times}10^{-4}$ | 0.7 | 1 | $9.94{\times}10^{4}$ | 23.4% | $9.44{\times}10^{-5}$ | 89.2% | $1.00{\times}10^{-4}$ | 96.6% | $1.00{\times}10^{-4}$ | 95.2% |
| 9 | 10000 | 30 | 100 | 23 | $1{\times}10^{-5}$ | 0.7 | 1 | $9.92{\times}10^{4}$ | 23.5% | $9.73{\times}10^{-5}$ | 94.7% | $9.99{\times}10^{-6}$ | 95.3% | $1.00{\times}10^{-5}$ | 94.7% |
| 10 | 10000 | 30 | 250 | 17.7 | $1{\times}10^{-4}$ | 0.7 | 1 | $5.07{\times}10^{4}$ | 14.8% | $9.86{\times}10^{-5}$ | 94.0% | $1.01{\times}10^{-4}$ | 95.2% | $1.00{\times}10^{-4}$ | 94.7% |
| 11 | 10000 | 30 | 250 | 20 | $1{\times}10^{-4}$ | 0.7 | 1 | $1.01{\times}10^{5}$ | 14.9% | $9.79{\times}10^{-5}$ | 93.5% | $1.00{\times}10^{-4}$ | 95.5% | $1.00{\times}10^{-4}$ | 94.6% |
| 12 | 10000 | 30 | 500 | 17.7 | $1{\times}10^{-4}$ | 0.7 | 1 | $1.01{\times}10^{5}$ | 10.5% | $9.93{\times}10^{-5}$ | 94.6% | $1.00{\times}10^{-4}$ | 95.4% | $1.00{\times}10^{-4}$ | 95.0% |
| 13 | 10000 | 30 | 250 | 53 | $1{\times}10^{-4}$ | 0.9 | 1 | $5.03{\times}10^{4}$ | 27.1% | $9.30{\times}10^{-5}$ | 88.4% | $1.00{\times}10^{-4}$ | 96.7% | $1.00{\times}10^{-4}$ | 94.9% |
| 14 | 10000 | 30 | 500 | 46.1 | $1{\times}10^{-4}$ | 0.9 | 1 | $5.04{\times}10^{4}$ | 19.2% | $9.66{\times}10^{-5}$ | 93.3% | $1.00{\times}10^{-4}$ | 95.9% | $1.00{\times}10^{-4}$ | 95.0% |
| 15 | 10000 | 30 | 500 | 53 | $1{\times}10^{-4}$ | 0.9 | 1 | $1.01{\times}10^{5}$ | 19.3% | $9.61{\times}10^{-5}$ | 91.2% | $1.00{\times}10^{-4}$ | 96.4% | $1.00{\times}10^{-4}$ | 94.9% |
| 16 | 10000 | 30 | 500 | 53 | $1{\times}10^{-5}$ | 0.9 | 1 | $1.00{\times}10^{5}$ | 19.2% | $9.82{\times}10^{-6}$ | 94.6% | $1.00{\times}10^{-5}$ | 95.4% | $1.01{\times}10^{-5}$ | 95.0% |
| 17 | 10000 | 30 | 1000 | 46.1 | $1{\times}10^{-4}$ | 0.9 | 1 | $1.01{\times}10^{5}$ | 13.5% | $9.80{\times}10^{-5}$ | 93.6% | $1.00{\times}10^{-4}$ | 95.5% | $1.00{\times}10^{-4}$ | 94.8% |
| 18 | 10000 | 30 | 1000 | 55.2 | $1{\times}10^{-4}$ | 0.9 | 1 | $2.51{\times}10^{5}$ | 13.6% | $9.78{\times}10^{-5}$ | 92.7% | $1.00{\times}10^{-4}$ | 96.2% | $1.00{\times}10^{-4}$ | 95.1% |
| 19 | 10000 | 30 | 2000 | 39.1 | $1{\times}10^{-4}$ | 0.9 | 1 | $1.00{\times}10^{5}$ | 9.5% | $9.91{\times}10^{-5}$ | 94.8% | $1.00{\times}10^{-4}$ | 95.5% | $1.00{\times}10^{-4}$ | 95.0% |

**Table S4. Minimum sample size required to achieve the power ≥ 80% when comparing mutation rates of two datasets.**

The rows represent the average number of mutations per culture of the first strain ($m_1$) while the columns represent the fold difference between the number of mutations (assuming the same final culture size, i.e., $N_{t1} = N_{t2}$) for the first and the second strain ($m_2$ / $m_1$). The sample size is the same for both strains ($n = n_1 = n_2$).

**$\varepsilon = 1.0$**

| $m_1$ \ $m_2/m_1$ | **1.1** | **1.2** | **1.3** | **1.4** | **1.5** | **1.75** | **2** | **2.25** | **2.5** | **3** | **3.5** | **4** | **5** |
|---|---|---|---|---|---|---|---|---|---|---|---|---|---|
| **0.5** | 4083 | 1077 | 504 | 298 | 200 | 99 | 61 | 43 | 33 | 21 | 16 | 12 | 9 |
| **0.75** | 2954 | 781 | 366 | 217 | 146 | 73 | 45 | 32 | 24 | 16 | 12 | 10 | 7 |
| **1** | 2378 | 630 | 296 | 176 | 118 | 59 | 37 | 26 | 20 | 13 | 10 | 8 | 6 |
| **1.25** | 2025 | 537 | 253 | 150 | 101 | 51 | 32 | 23 | 17 | 12 | 9 | 7 | 5 |
| **1.5** | 1784 | 474 | 223 | 133 | 90 | 45 | 28 | 20 | 16 | 11 | 8 | 6 | 5 |
| **1.75** | 1608 | 428 | 202 | 120 | 81 | 41 | 26 | 19 | 14 | 10 | 7 | 6 | 4 |
| **2** | 1473 | 392 | 185 | 110 | 75 | 38 | 24 | 17 | 13 | 9 | 7 | 6 | 4 |
| **2.5** | 1279 | 341 | 161 | 96 | 65 | 33 | 21 | 15 | 12 | 8 | 6 | 5 | 4 |
| **3** | 1144 | 305 | 144 | 86 | 58 | 30 | 19 | 14 | 11 | 7 | 6 | 5 | 4 |
| **3.5** | 1043 | 279 | 132 | 79 | 54 | 27 | 18 | 13 | 10 | 7 | 5 | 4 | 3 |
| **4** | 965 | 258 | 122 | 73 | 50 | 25 | 16 | 12 | 9 | 7 | 5 | 4 | 3 |
| **4.5** | 903 | 241 | 115 | 69 | 47 | 24 | 15 | 11 | 9 | 6 | 5 | 4 | 3 |
| **5** | 851 | 228 | 108 | 65 | 44 | 23 | 15 | 11 | 8 | 6 | 5 | 4 | 3 |
| **6** | 770 | 206 | 98 | 59 | 40 | 21 | 13 | 10 | 8 | 6 | 4 | 4 | 3 |
| **7** | 710 | 190 | 91 | 54 | 37 | 19 | 13 | 9 | 7 | 5 | 4 | 4 | 3 |
| **8** | 662 | 178 | 85 | 51 | 35 | 18 | 12 | 9 | 7 | 5 | 4 | 3 | 3 |
| **9** | 623 | 167 | 80 | 48 | 33 | 17 | 11 | 8 | 7 | 5 | 4 | 3 | 3 |
| **10** | 591 | 159 | 76 | 46 | 31 | 16 | 11 | 8 | 6 | 5 | 4 | 3 | 3 |
| **12.5** | 530 | 142 | 68 | 41 | 28 | 15 | 10 | 7 | 6 | 4 | 4 | 3 | 3 |
| **15** | 485 | 131 | 63 | 38 | 26 | 14 | 9 | 7 | 6 | 4 | 3 | 3 | 2 |
| **17.5** | 452 | 122 | 58 | 35 | 24 | 13 | 9 | 7 | 5 | 4 | 3 | 3 | 2 |
| **20** | 425 | 115 | 55 | 33 | 23 | 12 | 8 | 6 | 5 | 4 | 3 | 3 | 2 |
| **22.5** | 403 | 109 | 52 | 32 | 22 | 12 | 8 | 6 | 5 | 4 | 3 | 3 | 2 |
| **25** | 385 | 104 | 50 | 30 | 21 | 11 | 8 | 6 | 5 | 4 | 3 | 3 | 2 |
| **27.5** | 369 | 100 | 48 | 29 | 20 | 11 | 7 | 6 | 5 | 4 | 3 | 3 | 2 |
| **30** | 355 | 96 | 46 | 28 | 20 | 11 | 7 | 6 | 5 | 3 | 3 | 3 | 2 |
| **35** | 333 | 90 | 44 | 27 | 18 | 10 | 7 | 5 | 4 | 3 | 3 | 3 | 2 |
| **40** | 315 | 85 | 41 | 25 | 18 | 10 | 7 | 5 | 4 | 3 | 3 | 3 | 2 |
| **45** | 300 | 82 | 39 | 24 | 17 | 9 | 6 | 5 | 4 | 3 | 3 | 2 | 2 |
| **50** | 288 | 78 | 38 | 23 | 16 | 9 | 6 | 5 | 4 | 3 | 3 | 2 | 2 |
| **60** | 268 | 73 | 35 | 22 | 15 | 8 | 6 | 5 | 4 | 3 | 3 | 2 | 2 |
| **70** | 253 | 69 | 33 | 21 | 14 | 8 | 6 | 4 | 4 | 3 | 3 | 2 | 2 |
| **80** | 240 | 65 | 32 | 20 | 14 | 8 | 5 | 4 | 4 | 3 | 3 | 2 | 2 |
| **90** | 227 | 62 | 31 | 19 | 13 | 8 | 5 | 4 | 4 | 3 | 3 | 2 | 2 |
| **100** | 221 | 60 | 30 | 18 | 13 | 7 | 5 | 4 | 4 | 3 | 2 | 2 | 2 |

**$\varepsilon = 0.5$**

| $m_2/m_1$ \ $m_1$ | **1.1** | **1.2** | **1.3** | **1.4** | **1.5** | **1.75** | **2** | **2.25** | **2.5** | **3** | **3.5** | **4** | **5** |
|---|---|---|---|---|---|---|---|---|---|---|---|---|---|
| **0.5** | 5537 | 1458 | 680 | 401 | 268 | 132 | 82 | 57 | 43 | 28 | 20 | 16 | 11 |
| **0.75** | 3921 | 1034 | 484 | 285 | 191 | 95 | 59 | 41 | 31 | 20 | 15 | 12 | 8 |
| **1** | 3101 | 819 | 384 | 227 | 152 | 76 | 47 | 33 | 25 | 17 | 12 | 10 | 7 |
| **1.25** | 2601 | 688 | 323 | 191 | 128 | 64 | 40 | 28 | 21 | 14 | 11 | 8 | 6 |
| **1.5** | 2263 | 599 | 281 | 167 | 112 | 56 | 35 | 25 | 19 | 13 | 9 | 8 | 5 |
| **1.75** | 2018 | 535 | 251 | 149 | 100 | 50 | 32 | 22 | 17 | 11 | 9 | 7 | 5 |
| **2** | 1831 | 485 | 228 | 136 | 91 | 46 | 29 | 20 | 16 | 11 | 8 | 6 | 5 |
| **2.5** | 1563 | 415 | 195 | 116 | 78 | 39 | 25 | 18 | 14 | 9 | 7 | 6 | 4 |
| **3** | 1380 | 367 | 173 | 103 | 70 | 35 | 22 | 16 | 12 | 8 | 6 | 5 | 4 |
| **3.5** | 1245 | 331 | 156 | 93 | 63 | 32 | 20 | 15 | 11 | 8 | 6 | 5 | 4 |
| **4** | 1141 | 304 | 143 | 86 | 58 | 29 | 19 | 13 | 10 | 7 | 6 | 5 | 4 |
| **4.5** | 1059 | 282 | 133 | 80 | 54 | 27 | 18 | 13 | 10 | 7 | 5 | 4 | 3 |
| **5** | 991 | 264 | 125 | 75 | 51 | 26 | 17 | 12 | 9 | 6 | 5 | 4 | 3 |
| **6** | 886 | 237 | 112 | 67 | 46 | 23 | 15 | 11 | 9 | 6 | 5 | 4 | 3 |
| **7** | 808 | 216 | 102 | 61 | 42 | 21 | 14 | 10 | 8 | 6 | 4 | 4 | 3 |
| **8** | 748 | 200 | 95 | 57 | 39 | 20 | 13 | 9 | 7 | 5 | 4 | 4 | 3 |
| **9** | 699 | 187 | 89 | 53 | 36 | 19 | 12 | 9 | 7 | 5 | 4 | 3 | 3 |
| **10** | 659 | 177 | 84 | 50 | 34 | 18 | 12 | 9 | 7 | 5 | 4 | 3 | 3 |
| **12.5** | 584 | 157 | 75 | 45 | 31 | 16 | 10 | 8 | 6 | 5 | 4 | 3 | 3 |
| **15** | 530 | 142 | 68 | 41 | 28 | 15 | 10 | 7 | 6 | 4 | 3 | 3 | 3 |
| **17.5** | 490 | 132 | 63 | 38 | 26 | 14 | 9 | 7 | 5 | 4 | 3 | 3 | 2 |
| **20** | 458 | 123 | 59 | 36 | 25 | 13 | 9 | 6 | 5 | 4 | 3 | 3 | 2 |
| **22.5** | 432 | 116 | 56 | 34 | 23 | 12 | 8 | 6 | 5 | 4 | 3 | 3 | 2 |
| **25** | 411 | 111 | 53 | 32 | 22 | 12 | 8 | 6 | 5 | 4 | 3 | 3 | 2 |
| **27.5** | 393 | 106 | 51 | 31 | 21 | 11 | 8 | 6 | 5 | 4 | 3 | 3 | 2 |
| **30** | 377 | 102 | 49 | 30 | 21 | 11 | 7 | 6 | 5 | 4 | 3 | 3 | 2 |
| **35** | 351 | 95 | 46 | 28 | 19 | 10 | 7 | 5 | 4 | 3 | 3 | 3 | 2 |
| **40** | 331 | 90 | 43 | 26 | 18 | 10 | 7 | 5 | 4 | 3 | 3 | 3 | 2 |
| **45** | 314 | 85 | 41 | 25 | 17 | 10 | 7 | 5 | 4 | 3 | 3 | 2 | 2 |
| **50** | 300 | 81 | 39 | 24 | 17 | 9 | 6 | 5 | 4 | 3 | 3 | 2 | 2 |
| **60** | 278 | 76 | 37 | 22 | 16 | 9 | 6 | 5 | 4 | 3 | 3 | 2 | 2 |
| **70** | 261 | 71 | 34 | 21 | 15 | 8 | 6 | 5 | 4 | 3 | 3 | 2 | 2 |
| **80** | 247 | 67 | 33 | 20 | 14 | 8 | 6 | 4 | 4 | 3 | 3 | 2 | 2 |
| **90** | 236 | 64 | 31 | 19 | 14 | 8 | 5 | 4 | 4 | 3 | 3 | 2 | 2 |
| **100** | 227 | 62 | 30 | 19 | 13 | 7 | 5 | 4 | 4 | 3 | 2 | 2 | 2 |

$\varepsilon = 0.1$

| $m_2/m_1$ \ $m_1$ | **1.1** | **1.2** | **1.3** | **1.4** | **1.5** | **1.75** | **2** | **2.25** | **2.5** | **3** | **3.5** | **4** | **5** |
|---|---|---|---|---|---|---|---|---|---|---|---|---|---|
| **0.5** | 13673 | 3586 | 1668 | 980 | 653 | 319 | 196 | 136 | 101 | 65 | 47 | 36 | 24 |
| **0.75** | 9351 | 2455 | 1143 | 672 | 448 | 220 | 135 | 94 | 70 | 45 | 33 | 25 | 17 |
| **1** | 7178 | 1886 | 879 | 517 | 345 | 169 | 104 | 73 | 54 | 35 | 25 | 20 | 14 |
| **1.25** | 5868 | 1543 | 719 | 423 | 283 | 139 | 86 | 60 | 45 | 29 | 21 | 16 | 11 |
| **1.5** | 4989 | 1312 | 612 | 361 | 241 | 119 | 73 | 51 | 38 | 25 | 18 | 14 | 10 |
| **1.75** | 4357 | 1147 | 535 | 315 | 211 | 104 | 64 | 45 | 34 | 22 | 16 | 13 | 9 |
| **2** | 3881 | 1022 | 477 | 281 | 188 | 93 | 58 | 40 | 30 | 20 | 14 | 11 | 8 |
| **2.5** | 3208 | 845 | 395 | 233 | 156 | 77 | 48 | 34 | 25 | 17 | 12 | 10 | 7 |
| **3** | 2753 | 726 | 340 | 201 | 135 | 67 | 41 | 29 | 22 | 15 | 11 | 8 | 6 |
| **3.5** | 2425 | 640 | 300 | 177 | 119 | 59 | 37 | 26 | 20 | 13 | 10 | 8 | 5 |
| **4** | 2176 | 575 | 269 | 159 | 107 | 53 | 33 | 23 | 18 | 12 | 9 | 7 | 5 |
| **4.5** | 1981 | 523 | 245 | 145 | 98 | 49 | 30 | 21 | 16 | 11 | 8 | 6 | 5 |
| **5** | 1822 | 482 | 226 | 134 | 90 | 45 | 28 | 20 | 15 | 10 | 8 | 6 | 4 |
| **6** | 1582 | 419 | 196 | 116 | 78 | 39 | 25 | 17 | 13 | 9 | 7 | 5 | 4 |
| **7** | 1406 | 373 | 175 | 104 | 70 | 35 | 22 | 16 | 12 | 8 | 6 | 5 | 4 |
| **8** | 1273 | 337 | 159 | 94 | 63 | 32 | 20 | 14 | 11 | 7 | 6 | 5 | 4 |
| **9** | 1167 | 310 | 146 | 87 | 58 | 29 | 19 | 13 | 10 | 7 | 5 | 4 | 3 |
| **10** | 1081 | 287 | 135 | 80 | 54 | 27 | 17 | 12 | 10 | 7 | 5 | 4 | 3 |
| **12.5** | 923 | 245 | 116 | 69 | 47 | 24 | 15 | 11 | 8 | 6 | 5 | 4 | 3 |
| **15** | 814 | 217 | 102 | 61 | 41 | 21 | 13 | 10 | 8 | 5 | 4 | 4 | 3 |
| **17.5** | 734 | 196 | 92 | 55 | 37 | 19 | 12 | 9 | 7 | 5 | 4 | 3 | 3 |
| **20** | 673 | 179 | 85 | 51 | 34 | 18 | 11 | 8 | 7 | 5 | 4 | 3 | 3 |
| **22.5** | 624 | 166 | 79 | 47 | 32 | 17 | 11 | 8 | 6 | 4 | 4 | 3 | 3 |
| **25** | 583 | 156 | 74 | 44 | 30 | 16 | 10 | 7 | 6 | 4 | 3 | 3 | 2 |
| **27.5** | 550 | 147 | 70 | 42 | 29 | 15 | 10 | 7 | 6 | 4 | 3 | 3 | 2 |
| **30** | 521 | 139 | 66 | 40 | 27 | 14 | 9 | 7 | 5 | 4 | 3 | 3 | 2 |
| **35** | 475 | 127 | 61 | 36 | 25 | 13 | 9 | 6 | 5 | 4 | 3 | 3 | 2 |
| **40** | 440 | 118 | 56 | 34 | 23 | 12 | 8 | 6 | 5 | 4 | 3 | 3 | 2 |
| **45** | 411 | 110 | 53 | 32 | 22 | 12 | 8 | 6 | 5 | 4 | 3 | 3 | 2 |
| **50** | 388 | 104 | 50 | 30 | 21 | 11 | 7 | 6 | 5 | 3 | 3 | 3 | 2 |
| **60** | 351 | 94 | 45 | 27 | 19 | 10 | 7 | 5 | 4 | 3 | 3 | 2 | 2 |
| **70** | 324 | 87 | 42 | 25 | 18 | 9 | 6 | 5 | 4 | 3 | 3 | 2 | 2 |
| **80** | 302 | 82 | 39 | 24 | 17 | 9 | 6 | 5 | 4 | 3 | 3 | 2 | 2 |
| **90** | 285 | 77 | 37 | 23 | 16 | 9 | 6 | 5 | 4 | 3 | 3 | 2 | 2 |
| **100** | 271 | 73 | 35 | 22 | 15 | 8 | 6 | 4 | 4 | 3 | 3 | 2 | 2 |

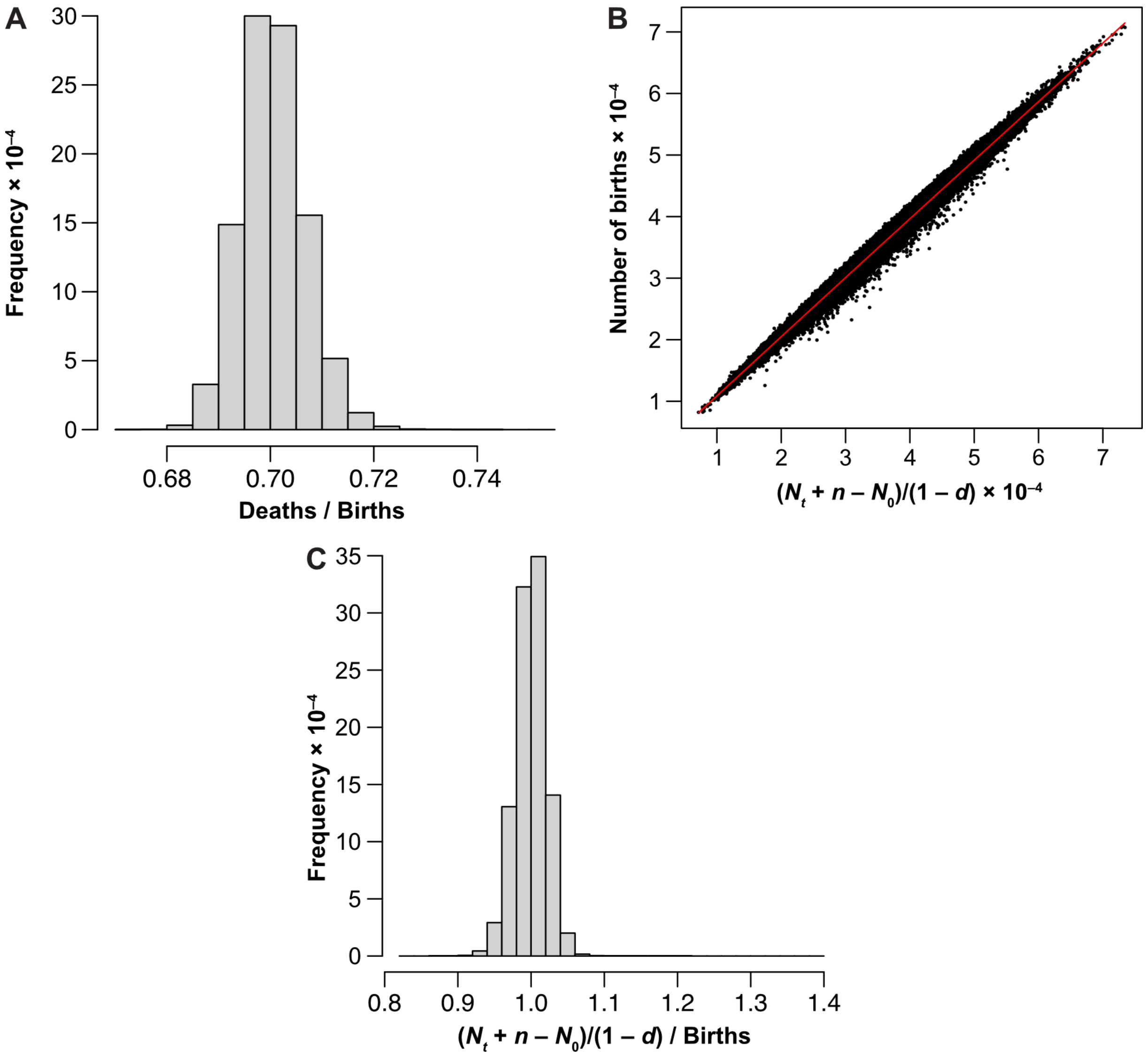


**Figure S1. The correlation between the number of births and final culture size for data used in Line 5 of Table 6.**

**(A)** The histogram depicts the observed proportion $d$ = Deaths / Births from $10^6$ simulated test tubes. The nominal value is 0.7, the mean is 0.700, and standard deviation equals 0.00622. **(B)** The scatter plot shows the correlation between $(N_t + n - N_0)/(1 - d)$, the estimate of the number of cell divisions used in calculations of $\mu$, and the true number of wild-type births from $10^6$ simulated test tubes. Each black dot represents a single test tube. A trend line (red) has been added for reference. The Pearson's correlation coefficient is 0.997. **(C)** The histogram uses the same data as in (B), but expresses $(N_t + n - N_0)/(1 - d)$ as a fraction of the true number of births. The minimum value is 0.833, the maximum is 1.24, the mean is 1.00, and the standard deviation is 0.0212.
